# The beryllium fluoride form of the Na^+^,K^+^-ATPase and the intermediates of the functional cycle

**DOI:** 10.1101/2021.11.08.467847

**Authors:** Marlene U. Fruergaard, Ingrid Dach, Jacob L. Andersen, Mette Ozol, Azadeh Shasavar, Esben M. Quistgaard, Hanne Poulsen, Natalya U. Fedosova, Poul Nissen

## Abstract

The Na^+^,K^+^-ATPase generates electrochemical gradients of Na^+^ and K^+^ across the plasma membrane. Here, we describe a 4.0 Å resolution crystal structure of the pig kidney Na^+^,K^+^-ATPase stabilized by beryllium fluoride (denoted E2-BeF_x_). The structure shows high resemblance to the E2P phosphoenzyme obtained by phosphorylation from inorganic phosphate (P_i_) and stabilised by cardiotonic steroids, and reveals a Mg^2+^ bound near the ion binding site II. Anomalous Fourier analysis of the crystals soaked in Rb^+^ (K^+^ congener) followed by a low resolution rigid-body refinement (6.9-7.5 Å) revealed pre-occlusion transitions leading to activation of the desphosphorylation reaction. Mg^2+^ location indicates a site of an initial K^+^ recognition and acceptance upon binding to the outward-open E2P state after Na^+^ release. Despite the overall structural resemblance to the P_i_-induced E2P phosphoform, BeF_x_ inhibited enzyme is able to bind both ADP/ATP and ions - features that relate E2-BeF_x_ complex to an intermediate of the functional cycle of the Na^+^,K^+^-ATPase prior E2P.

## Introduction

Na^+^,K^+^-ATPase maintains physiological concentrations and gradients of Na^+^ and K^+^, which are crucial for animal cells. The enzyme is a binary complex of a large α-subunit of the P-type ATPase family responsible for ion transport and enzymatic reaction, and a β-subunit acting as a chaperone and functional modulator. Furthermore, regulatory FXYD proteins may associate with the complex in a tissue-dependent manner. The ion exchange is driven by ATP hydrolysis via enzyme autophosphorylation and dephosphorylation, and a minimal scheme of the Na^+^,K^+^-ATPase cycle, also termed the Post-Albers scheme, includes two major conformations, E1 and E2, in their phosphorylated and non-phosphorylated states (1,2). These four states display different ion binding affinities and provide access to the ion sites either from inward/cytoplasmic or outward/extracellular sides of the membrane. The dephosphorylation of the phosphoenzyme pool (preformed by Na^+^-dependent phosphorylation) in response to addition of either ADP or K^+^ was shown to be biphasic in both cases, in agreement with the presence of two phosphointermediates in the reaction cycle (E1P and E2P). The amplitude of the fast phase of P_i_ release in ADP- or K^+^- chase experiments reflected respective fractions of ADP-sensitive (E1P) and K^+^-sensitive (E2P) phosphoenzymes in the Post-Albers scheme. However, the sum of these two phosphointermediates considerably exceeded the amount of initially phosphorylated enzyme (3), exposing insufficiency of a two-state model (E1P/E2P) for phosphointermediates. The number and the nature of the phosphoenzyme conformations have therefore been debated, and a consensus was found in the existence of three phosphoenzymes: ADP-sensitive K^+^-insensitive (E1P), ADP- and K^+^-sensitive (E*P), and ADP-insensitive K^+^-sensitive (E2P) phosphoenzymes (3-5). Similar dephosphorylation phenomena were also observed for the closely related sarco/endoplasmic reticulum Ca^2+^-ATPase (SERCA) (4).

Recent success in crystallization revealed structures mimicking several native phosphointermediates of the Na^+^,K^+^-ATPase. Protein complexes with metal fluorides (i.e. beryllium, aluminium, and magnesium fluorides, denoted BeF_x_, AlF_x_ and MgF_x_ due to varying degrees of water coordination and therefore different fluoride stoichiometries and net charges) mimic different states of phosphoryl transfer reactions and phosphointermediates. Thus, a MgF_x_-complex resembles an [K_2_]E2·P_i_ state with occluded K^+^ and non-covalently bound phosphate (P_i_), while E1·AlF_4_^−^·ADP contains three occluded Na^+^ ions and represents an intermediate leading to the [Na_3_]E1P-ADP phosphoenzyme (6,7). The BeF_x_-complex of the Na^+^,K^+^-ATPase (8,9) is structurally similar to the P_i_-induced (“backdoor” phosphorylated) E2P phosphoenzyme stabilised by cardiotonic steroids, the only structure so far of a genuine phosphoenzyme (10,11).

The flexibility and intermediary functional properties of the BeF_x_ complex were anticipated earlier from the presence of a characteristic H^+^ leak current mediated by Na^+^,K^+^-ATPase (12) and in SERCA from the measurements of Ca^2+^-occlusion (13) and Ca^2+^ mediated reactivation (14).

Here we describe functional properties of the E2-BeF_x_ complex of Na^+^,K^+^-ATPase and structural re-arrangements in its crystal structure induced by binding of Rb^+^ as a K^+^ congener. Close structural resemblance to the P_i_-induced E2P phosphoenzyme stabilised by cardiotonic steroids (10,11) includes outward-open ion binding sites, which prompted investigations of the effect of extracellular Mg^2+^.

Notably, we also find that BeF_x_ dissociation from the enzyme is accelerated by both nucleotide and ion binding – these are characteristics that relate the E2-BeF_x_ structure to the ADP- and K^+^-sensitive (E*P) phosphoenzyme.

## Results

### Formation of the BeF_x_ complex

The BeF_x_ inhibited complex of Na^+^,K^+^-ATPase was formed by pre-incubation of the membraneous enzyme in 20 mM histidine pH 7.0, 10 mM NaF, 0.5 mM MgCl_2_, 20 μM BeSO_4_ and used in all biochemical studies as well as initial material for crystallization.

### Overall crystal structure of the BeF_x_-complex of the Na^+^,K^+^-ATPase

The crystal structure of pig kidney Na^+^,K^+^-ATPase (α_1_, β_1_ and FXYD2 also known as γ) was determined by molecular replacement using the ouabain-bound P_i_-induced (E2P-OBN) form as the starting model (PDB ID 4HYT) (10), and at later stage compared to the structure of a different crystal form of the E2-BeFx complex (PDB 7D91 (9)). The final model was refined against anisotropically truncated data extending to 4.0 Å, resulting in R_work_ and R_free_ values of 22.7 % and 27.6 %, respectively (Fig. 1A) (Table S1).

**Fig. 1.**
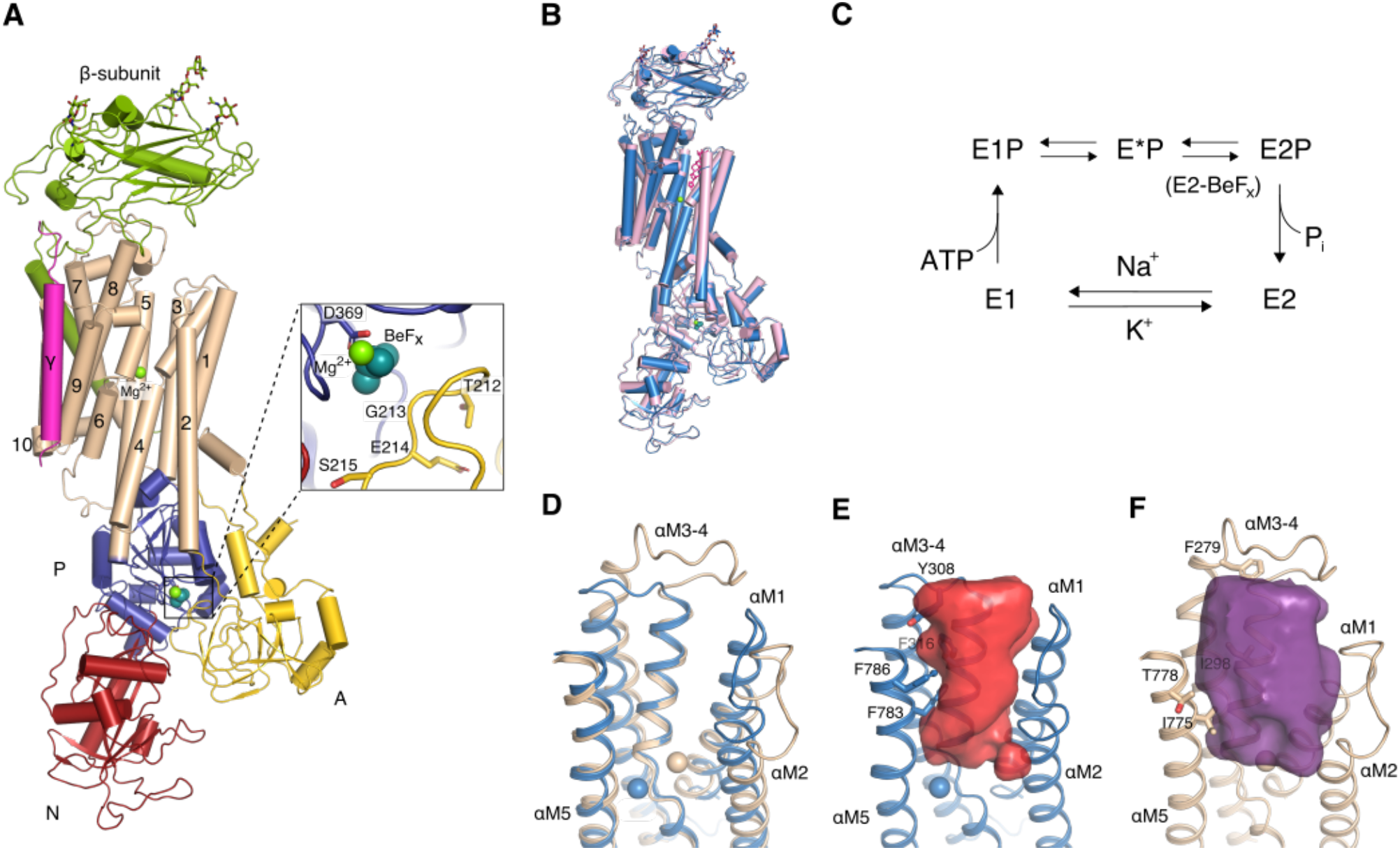
Crystal structure of E2-BeF_x_ state of Na^+^,K^+^-ATPase. (A) Cartoon representation of E2-BeF_x_ state colored according the different domains: red (nucleotide binding (N) domain), blue (phosphorylation (P) domain), yellow (activator (A) domain), wheat (transmembrane domain (TM) αM1-M10), green (β-subunit) and hotpink (γ-subunit). Close-up view of the phosphorylation site is shown in the *inset*. Mg^2+^ ions and the BeF_x_ bound to Asp369 are depicted as green and teal spheres, respectively. (B) Overall comparison of E2-BeF_x_ (blue) and P_i_-induced E2P (pink). Mg^2+^ ions are depicted as green spheres and ouabain is colored in hotpink. The structures were aligned on the transmembrane segment αM7-10. (C) The Post-Albers reaction scheme of Na, K-ATPase accumulating 3-pool model of phosphoenzymes. (D) Extracellular access channel of Na^+^,K^+^-ATPase E2-BeF_x_ (blue) aligned with SERCA E2-BeF_x_ (PDB ID 3B9B (14), wheat). The structures were aligned by αM5-M6. Bound Mg^2+^ ions are shown as spheres also in blue and wheat. (E,F) Water cavity representation of Na^+^,K^+^-ATPase E2-BeF_x_ (red) and SERCA E2-BeF_x_ (purple), showing a narrower entrance to the ion binding sites in the Na^+^,K^+^-ATPase E2-BeF_x_ complex. Cavities were calculated in HOLLOW(17).

BeF_x_ coordinated to the conserved Asp369 phosphorylation site in the conserved DKTG segment of the P domain was identified in unbiased, initial *F_o_*-*F_c_* difference map. Comparison shows similar arrangements of the P domains in both the P_i_-induced ouabain-bound E2P state (10) and in the present BeF_x_ structure. The TGES loop of the actuator (A) domain is in close proximity to BeF_x_ and protects the phosphorylation site from spontaneous hydrolysis (Fig. 1A). High structural similarity with E2P-OBN (root mean square deviations (rmsd) = 0.59 Å for all C_α_), also reveal subtle ouabain-induced rearrangements in the αM1-M4 segments of the earlier reported E2P-OBN structure. Slight outwards tilting of the αM1-M2 bundle is due to the extensive hydrogen-bonding network between the β-surface of ouabain and polar sidechains of αM1, αM2 and αM6 (10). Furthermore, the inhibitor-bound site makes the extracellular part of the αM3-M4 helices close in, causing a ~4° tilt of αM4 with pivot at Val322 (Fig. 1B) as compared to E2-BeF_x_.

The extracellular ion pathway is formed by the transmembrane αM1-M6 helices, which define an outward-open arrangement similar to sarcoplasmic reticulum Ca^2+^-ATPase (SERCA1a) in the Mg^2+^ stabilized E2-BeF_x_ form (14) (Fig. 1D). The arrangement of αM2-M5 helices in Na^+^,K^+^-ATPase, however, is slightly more compact and the residues lining the pathway have larger hydrophobic side chains (e.g. Tyr308, Phe316, Phe783 and Phe786 in Na^+^,K^+^-ATPase corresponding to Phe279, Ile298, Ile775 and Thr778 in SERCA) (Fig. 1E,F). The extracellular ion pathway is thus narrow but solvated, consistent with a low voltage sensitivity of K^+^ binding through such a pathway (15) and a voltage-insensitive release of the third and last Na^+^ ion from the Na^+^ bound state (16).

For the cation binding sites in the transmembrane domain, we observed a residual density near site II in the initial, unbiased F_o_-F_c_ difference map, similar to a Mg^2+^ site in the E2P-OBN complex (10). Indeed, the E2-BeF_x_ crystals were grown in the presence of 175 mM MgCl_2_, and Mg^2+^ binding is likely to stabilize an open conformation as seen also for the E2-BeF_x_ complex of SERCA1a (14), although the site is shifted (Fig. 1D). The Mg^2+^ site seems to overlaps with incoming K^+^ (see further below).

### P_i_-induced E2P and E2-BeF_x_ complexes have similar functional properties

Structural similarities between P_i_-induced E2P and E2-BeF_x_ complexes (Fig. 1B) are also supported by their functional properties, e.g. by interactions with cardiotonic steroids. Fig. 2 summarises kinetics of anthroylouabain binding to these complexes, which, judged from the crystallographic data, have very similar organization of the CTS binding cavities. The main conclusion from the kinetic experiments is that affinities of both forms to anthroylouabain are very high, with only small numeric differences in the values of association and dissociation rate constants. Although the effect of K^+^ on anthroylouabain affinity seems to be less pronounced, it is still in the direction of decreased affinity for both forms.

**Fig. 2.**
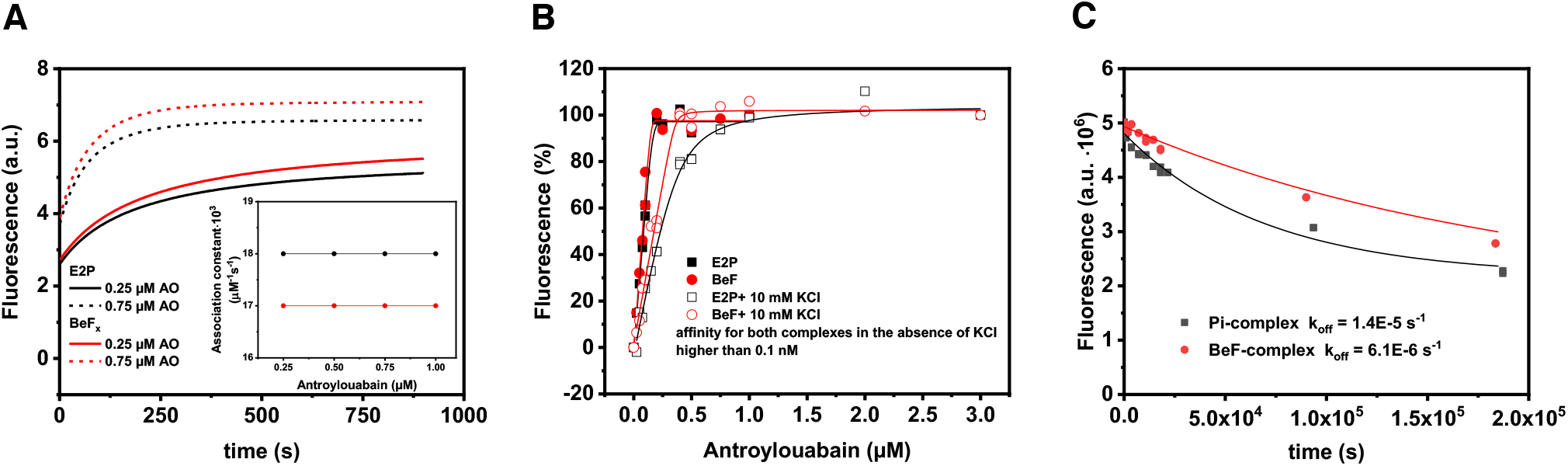
Interaction of P_i_-induced E2P and BeF_x_ complexes of Na^+^,K^+^-ATPase with anthroylouabain. Data obtained with P_i_-induced E2P are shown in black color, with BeF_x_ complex in red. (A) Changes in anthroylouabain (AO) fluorescence at two different concentrations, 0.25 μM (full line) and 0.75 μM (stippled line), were induced by its binding to the E2P and BeF_x_ complexes. Calculated second order association rate constants for the reactions are shown in the *inset.* (B) Amplitude of the fluorescence change as function of anthroylouabain concentration in the absence or presence of 10 mM KCl. (C) Time course of anthroylouabain dissociation from E2P and BeF_x_ complexes.

### Interactions of Na^+^, K^+^/Rb^+^ and Mg^2+^ with the E2-BeF_x_ complex

The E2-BeF_x_ crystal structure reveals that the cation binding cavity is open to the extracellular side and occupied by a Mg^2+^ ion. The homologous complex of SERCA in crystallized form also contained Mg^2+^ (14) (Fig. 1F), while in functional studies it was shown to bind and occlude Ca^2+^ (13). Do the ions bind to the E2-BeF_x_ complex of Na^+^,K^+^-ATPase, and what are the functional consequences of this binding? We investigated the interactions with Na^+^, K^+^ (Rb^+^as congener), and Mg^2+^.

#### Mg^2+^ binding does not affect Na^+^ and K^+^ binding

The effect of extracelluar Mg^2+^ on binding of Na^+^, K^+^ and H^+^ was investigated by two-electrode voltage-clamping on *Xenopus laevis* oocytes (a well-established model to study functional properties of the Na^+^,K^+^-ATPase), expressing human α_2_β_1_. In the absence of extracellular K^+^, the pump is distributed between the whole range of phosphoenzymes due to voltage-dependent binding and release of Na^+^. We assayed Na^+^ binding in the presence of 0 mM, 1 mM and 5 mM Mg^2+^ and found no difference in either apparent affinity for Na^+^ (Q/Vm curves) or the rate constants of release (Fig. S1A,B). We next examined the potency and efficacy of K^+^ binding by measuring the steady-state current at −30 mV in the presence of 115 mM Na^+^ with or without 5 mM Mg^2+^ at varying K^+^ concentrations. Again, there were no differences in currents suggesting that physiological concentrations of extracellular Mg^2+^ have no significant effects on binding of Na^+^ or K^+^ from the extracellular environment (Fig. S1C,D).

However, in the absence of extracellular Na^+^ and K^+^, an inwardly rectifying H^+^ leak current is observed for the Na^+^,K^+^-ATPase (7), in particular at hyperpolarized potentials. This inward leak, however, was significantly inhibited at hyperpolarized potentials by 5 mM or 20 mM Mg^2+^ (Fig. 3).

**Fig. 3.**
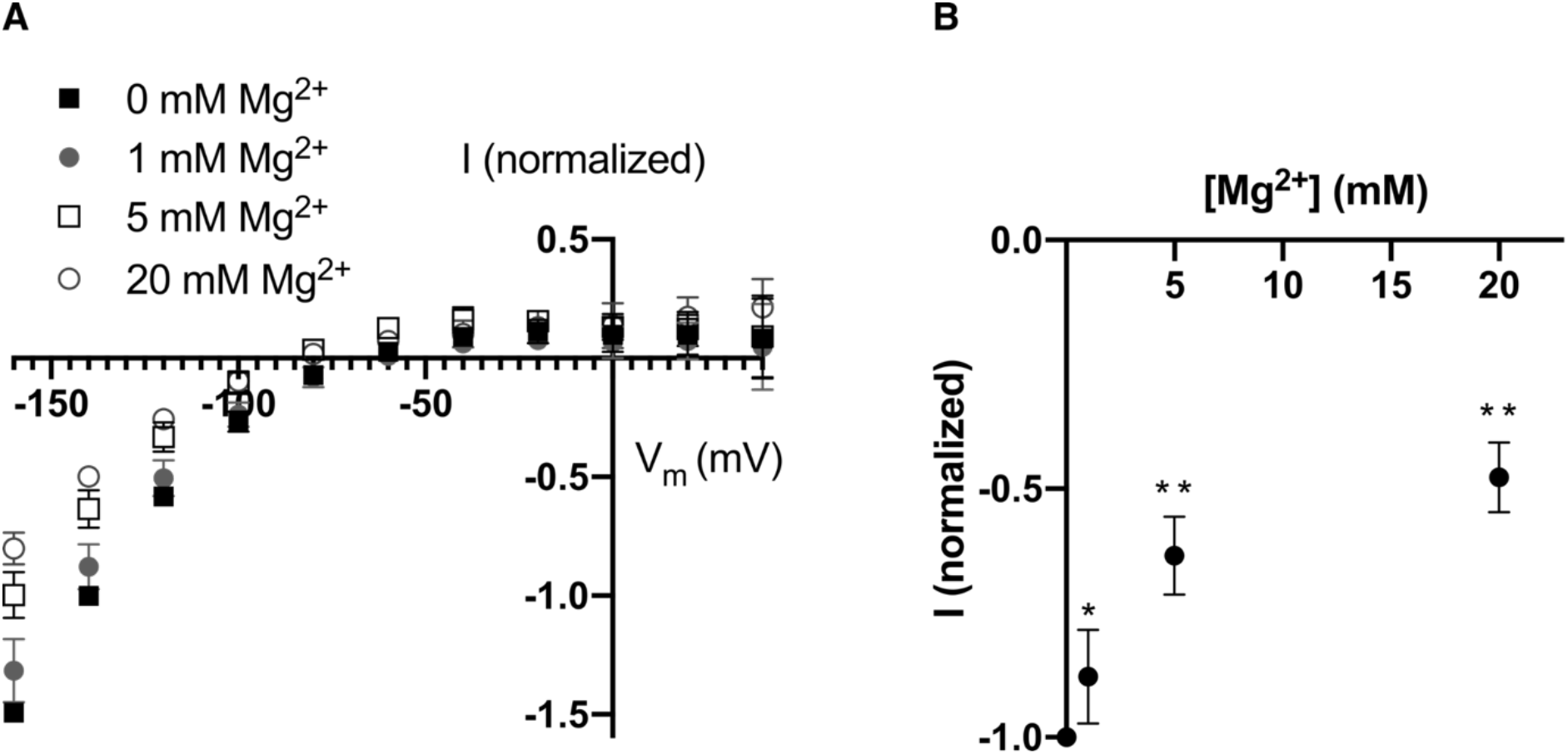
Inward leak current is affected by magnesium. (A) The ouabain-sensitive steady state current measured in the absence of extracellular sodium and potassium in response to membrane potential for different concentrations of magnesium. Current is normalized to −1 for 0 mM Mg^2+^ at −140 mV (n=4). (B) Leak currents at −140 mV. For 5 mM and 20 mM, the currents are significantly smaller (*, p<0.05; **, p<0.0001; Student’s t-test).

Thus, the affinity for the extracellular Mg^2+^ is so low that it does not interfere with Na^+^ or K^+^ binding under physiological conditions. Yet, when bound, it stabilizes the outward-open state and interferes with a H^+^ leak current.

### Beryllium fluoride prevents oligomycin-induced Na^+^-occlusion

The E2-BeF_x_ form of SERCA1a is capable of Ca^2+^-occlusion (13). We therefore investigated Na^+^-occlusion by the E2-BeF_x_ form of the Na^+^,K^+^-ATPase, but failed to reveal any bound Na^+^ (Fig. 4A).

**Fig. 4.**
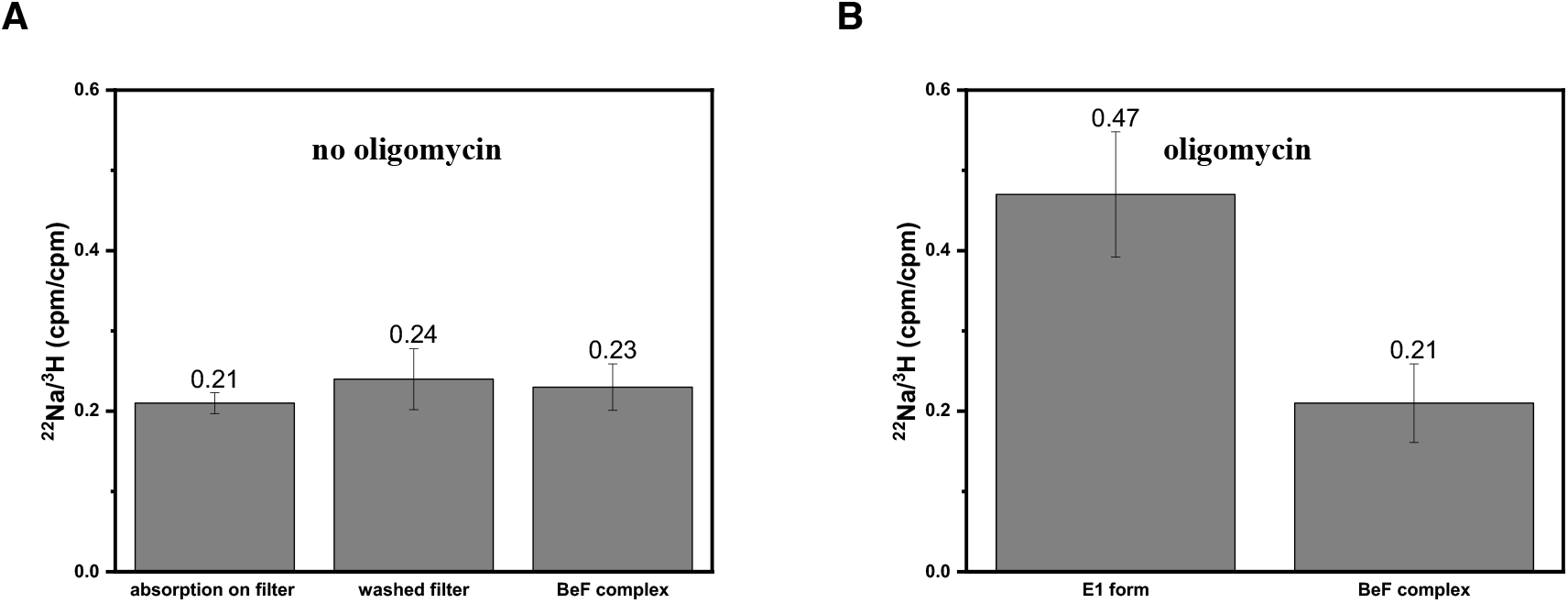
BeF_x_ binding prevents Na^+^-occlusion in the presence of oligomycin. Counts for ^22^Na^+^ on the filter are related to the counts for ^3^H^+^ used as internal standard for non-specific binding. The ^22^Na^+^/^3^H^+^ ratio (counts per filter) after filtration of 0.4 ml incubation media are shown for the samples of different compositions. (A) in the absence of added oligomycin: 1) no protein, non-specific binding without washing of the filter; 2) no protein, non-specific binding after washing of the filter; 3) Na^+^,K^+^-ATPase in the E2-BeF_x_ form, followed by washing of the filter. In the absence of Na^+^,K^+^-ATPase the ratio was not changed by washing, indicating that both isotopes interact with filter in the same way. (B) in the presence of oligomycin. 1) Na^+^,K^+^-ATPase in E1 conformation, followed by washing of the filter; 2) Na^+^,K^+^-ATPase in the E2-BeF_x_ form, followed by washing of the filter.

We repeated the experiment in the presence of oligomycin, which decreases the rate of Na^+^-release (18,19), and found an expected increase in ^22^Na^+^ bound to the E1 state, where the Na^+^,K^+^-ATPase occluded approximately 2.5 nmol per nmol protein (Fig. 4B). However, under the conditions of BeFx-complex formation the ^22^Na^+^/^3^H^+^ ratio was the same as in the absence of enzyme or absence of oligomycin (Fig.4A), implying that binding of BeF_x_ effectively prevented Na^+^-occlusion by the enzyme.

### K^+^-occlusion by the BeF_x_-complex

BeF_x_ binding to Na^+^,K^+^-ATPase is associated with an increase of RH421 fluorescence, similar to the response of this dye to enzyme phosphorylation, while addition of K^+^-ions induces a decrease in fluorescence (Fig. 5A, inset, and (20)). The values for both amlitude and k_obs_ for K-response extracted by fitting with a monoexponential function show a hyperbolic concentration dependence for the amplitude of fluorescence that decrease from its maximal level already at 2 mM KCl and a linear increase for k_obs_ (Fig. 5). Knowing, that K^+^ interacts with P_i_-induced E2P in the same way (21), we assumed direct binding of	K^+^ is followed by its occlusion in E2-BeF_x_ form. Indeed, direct measurements with ^86^Rb^+^ (as K^+^ congener) showed its accumulation in the E2-BeF_x_ form (20).

**Fig.5.**
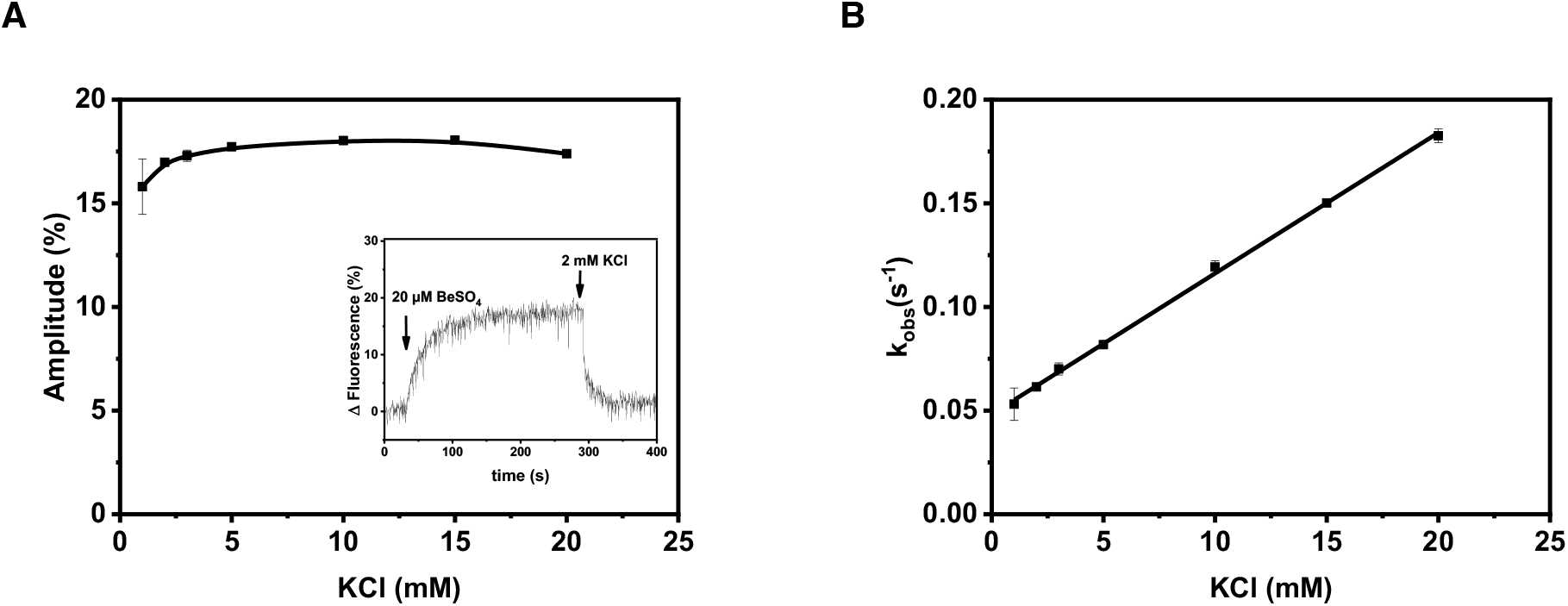
Interactions of K^+^ and BeF_x_-complex monitored by RH421 fluorescence. (A) The amplitude of the fluorescence change induced by addition of varying K^+^-concentration to pre-formed Na^+^,K^+^-ATPase-BeF_x_ complex. *Inset* illustrates changes in RH421fluorescence in response to addition of ligands to pig kidney enzyme. The changes of fluorescence are expressed as percentage of the initial level. (B) The observed rate constant of the fluorescence change (k_obs_) as function on K^+^-concentration. The values obtained from exponential fit are the means of three experiments ±S.E.

#### Rb^+^ binding and extracellular gate closure

Comparison of the E2-BeF_x_ (this study) and [Rb_2_]E2-MgF_x_ (6) states reveals the overall transitions of the extracellular gate closure upon K^+^/Rb^+^ binding. We tracked the binding process and mechanism of occlusion by crystal soaking procedures.

### Structural re-arrangements following Rb^+^-binding

#### Pre-occluded (Rb)E2-BeF_x_ states

Structural re-arrangements in the BeF_x_-complex due to Rb^+^ binding and occlusion were followed by soaking E2-BeF_x_ crystals. Two datasets were collected at 6.9 and 7.5 Å maximum resolution (consisting of 11,991 and 9,616 unique reflections, respectivley) at a wavelength of 0.814 Å with a strong anomalous signal for Rb^+^. They represent transition intermediates from the native E2-BeF_x_ with bound Mg^2+^ (Fig. 6A) towards the earlier reported occluded structure of [Rb_2_]E2-MgF_x_ (PDB 3KDP (6)). Indeed, the datasets for Rb^+^ soaked E2-BeF_x_ (here referred to as quick soak and long soak corresponding to 10 mM, 20 sec and 50 mM, 3 hours, respectively) produced strong anomalous difference peaks (5.8 to 8.4 σ) at the cation binding sites, reflecting exchange of Mg^2+^ with Rb^+^ (Fig. 6B,C,D). As the asymmetric unit of the P2_1_2_1_2_1_ crystals consists of two protomers, we have altogether four represenations of Rb^+^-soaked E2-BeF_x_ states.

**Fig. 6.**
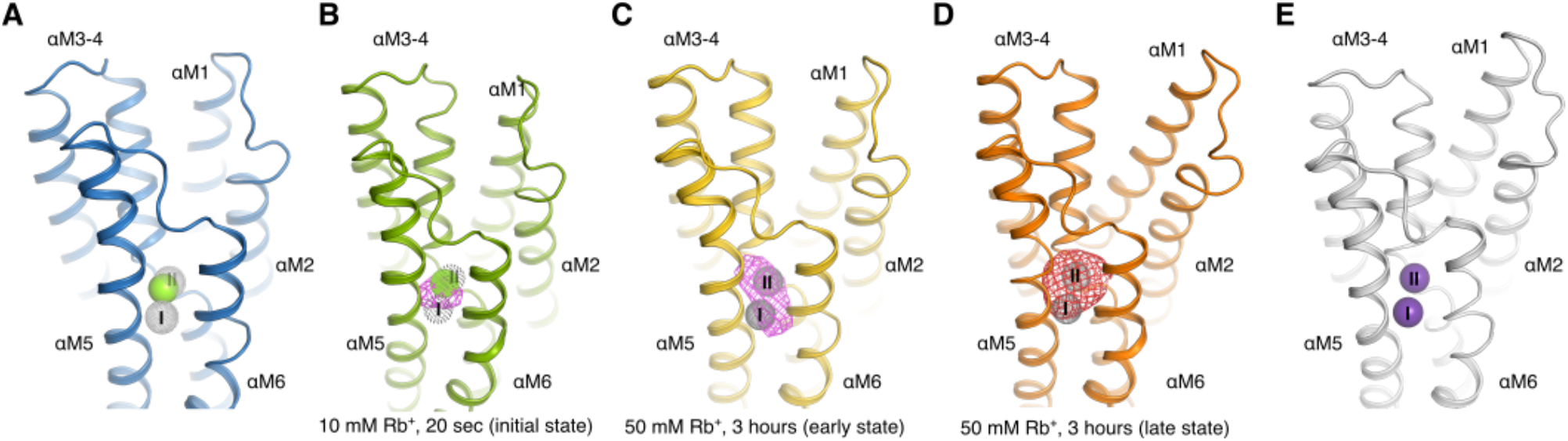
Structural re-arrangements following Rb^+^-binding to E2-BeF_x_. (A) Open E2-BeF_x_ Mg^2+^ bound form (this study). Mg^2+^ is shown as a green sphere, and the positions of site I and II are indicated by grey dotted spheres. (B) Initial binding form (10 mM Rb^+^, 20 sec). The 4-σ anomalous difference map is shown as a purple mesh, and cation binding sites are shown as in panel A. Mg^2+^ site from open E2-BeF_x_ state is shown as green dotted sphere. (C) Early (Rb)E2-BeF_x_ form (soaked with 50 mM for 3 hour, protomer 1). The 4-σ anomalous difference map is shown as a purple mesh, and cation binding sites are shown as in panel A. (D) Late (Rb)E2-BeF_x_ form (soaked with 50 mM for 3 hours, protomer 2). The anomalous difference map is contoured at 3σ (red mesh) and cation binding sites are shown as in panel A. (E) Occluded [Rb_2_]E2-MgF_x_ form (PDB ID 3KDP)(6). Rb^+^ ions are shown as purple spheres.

Quantitative determination of Rb^+^ occupancy was challenging due to the low resolution of the data sets, but occupancy analyses through Molecular Replacement with Single-wavelength Anomalous Diffraction (MR-SAD) refinement appeared consistent with two fully occupied binding sites for both protomers in the long Rb^+^-soak (occupancy >0.75). For the two quick soak conformations, one protomer was fully occupied, while the other showed partial occupancy (with an occupancy less than 0.1 for a second site, assuming a full occupancy of a first site). Interestingly, the anomalous difference peak for the partially occupied quick soak protomer overlapped with the Mg^2+^ site right in middle of the two K^+^ sites I and II), suggesting that initial Rb^+^/K^+^ binding takes place at this exposed site prior to formation of the properly coordinated K^+^ (or Rb^+^) sites (Fig. 6B). That said, the low resolution obviously precludes firm conclusions on this point.

Rigid-body model refinement for the cytosolic domains and individual transmembrane segments produced large improvements in crystallographic R-factors for model representations and converged robustly at conformational changes that were also consistent with unbiased omit map controls.

For all four protomers of the soaked crystals, the cytoplasmic domains exhibited weak density in the electron density maps, likely indicating flexibility (Fig. S2, S3). The E2-BeF_x_ MR model provided the relative position of the cytosolic headpiece based on some defined helices in each domain; thus, they are still modelled despite the weak density. The transmembrane helices, however, were clearly visible in the electron density map to model the C_α_ mainchain. The two protomers in the quick soak are similar to each other (rmsd = 1.16 Å, all C_α_; Fig. S2) and also show resemblance to protomer 1 in the long-soaked crystal (rmsd: ~1.27 Å, all C_α_). However, the last protomer (long soak protomer 2) has a different conformation for the transmembrane helices (rmsd = 1.96 Å between C_α_ of the two long-time soak protomers; Fig. S3). For simplicity, the protomers will therefore in the following be refered to as the initial (Rb)E2-BeF_x_ binding form (quick soak conformations), the early (Rb_2_)E2-BeF_x_ binding form (long time soak protomer 1) and the late (Rb_2_)E2-BeF_x_ binding form (long time soak protomer 2) when discussing structural changes.

#### Sequence of events

The refined atomic models for the E2-BeF_x_ form (this study) and [Rb_2_]E2-MgF_x_ form (6) represent the start and end points of the K^+^ binding, and the rigid body refined models of the soaked crystals, albeit determined at low resolution, provide valuable insights into the trajectories of the K^+^/Rb^+^ induced conformational changes that activate the dephosphorylation reaction.

Superposition of the Rb^+^-soaked structures and the native E2-BeF_x_ form based on the transmembrane αM7-M10 segment, revealed sequential closing movements of the M1-M4 helices (Fig. 7). K^+^ binding associates the extracellular part of αM4 to site II, most likely through mainchain carbonyls of Val322, Ala323 and Val325 engaging in coordination (6,22). In the initial binding (Rb)E2-BeF_x_ form, this results in αM4 tilting ~7° towards αM6 (Fig. 7A,B). In the early (Rb_2_)E2-BeF_x_ form, αM4 is further tilted ~5°, and the extracellular gate is closing. Through van der Waals contacts between αM1, αM2 and αM4, the αM1-M2 segment follows αM4 to an intermediate position via a ~2 Å translation towards the extracellular side (Fig. 7C,D)

**Fig. 7.**
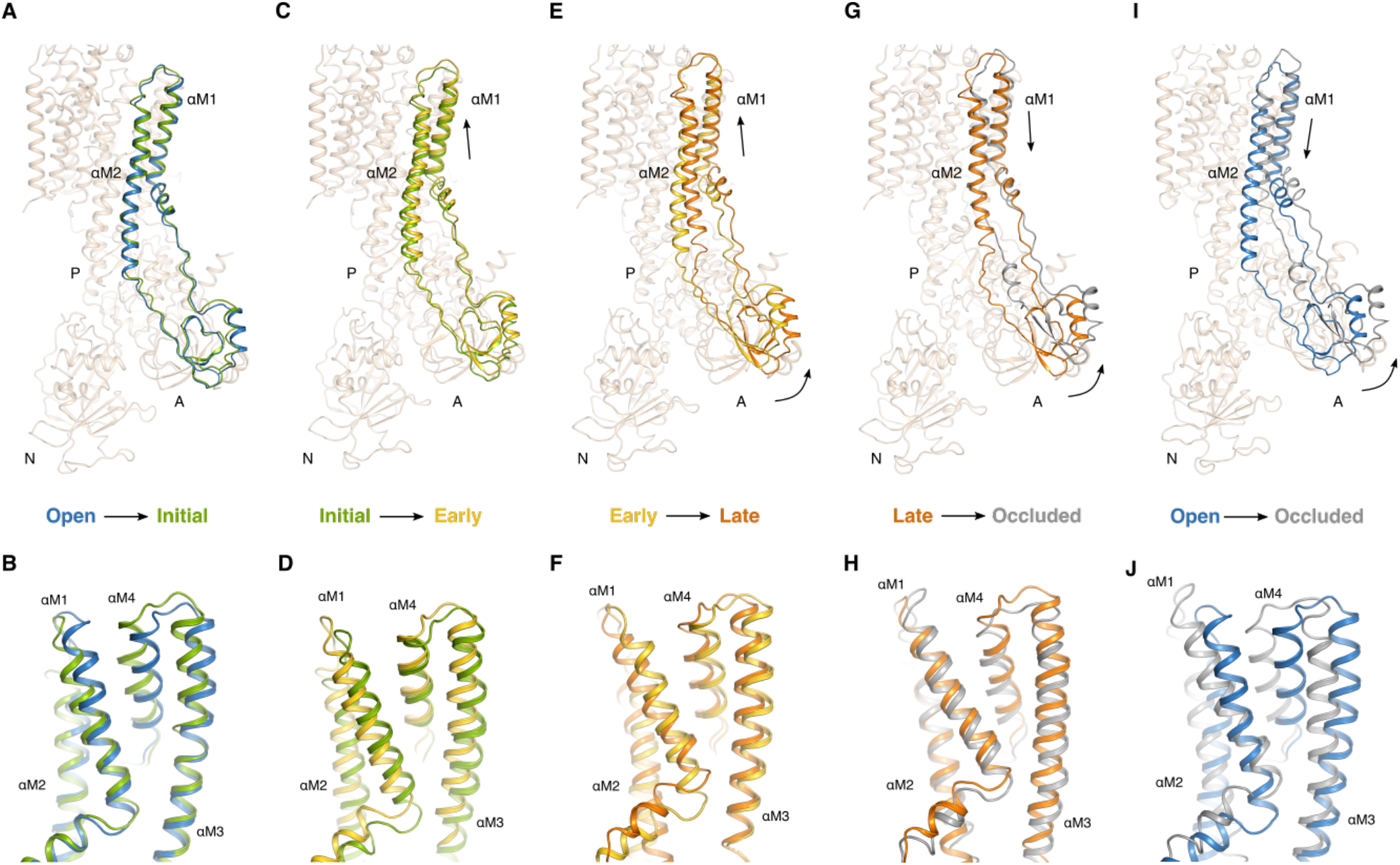
Sequential structural changes leading to occlusion of Rb^+^ in the E2 state. Structural differences (A and B) between the open E2-BeF_x_ Mg^2+^ (blue) and initial binding (Rb)E2-BeF_x_ form (green) in an overall view (A) and for the transmembrane membrane domain (B), respectively, (C and D) the initial (Rb)E2-BeF_x_ form (green) and the early (Rb)E2-BeF_x_ (gold), (E and F) Early (Rb)E2-BeF_x_ (gold) and late (Rb)E2-BeF_x_ form (orange), (G and H) late (Rb)E2-BeF_x_ form (orange) and fully occluded [Rb_2_]E2-MgFx (grey) (PDB ID 3KDP (6)), and (I and J) open E2-BeF_x_ Mg^2+^ (blue) and fully occluded [Rb_2_]E2-MgFx (grey). All structures were aligned on αM7-M10. For clarity, only the regions showing major conformational re-arrangements between the states have been highlighted.

In the next step, reaching the late (Rb_2_)E2-BeF_x_ form, the αM1-M2 helices are further translated towards to extracellular side (~4.5 Å) and ~6° lateral tilted towards the cytosolic side. The cytoplasmic part of the αM2 helix bends towards the A-αM3 linker region (Fig, 7E,F). The change of path is likely realized by a partial unwinding of the αM2 helix (M2 switch (23)), which also gains flexibility; as is indicated by poor density for the cytoplasmic end of αM2 (Fig. S3). The unwinding enables the A domain to rotate ~7 ° around the phosphorylation site of the P domain (towards the membrane). To complete the transition to the fully occluded [Rb_2_]E2-MgF_x_ complex (PDB ID 3KDP, (6)), the A domain must finish its rotation (~7 °), bringing the TGES motif into dephosphorylation mode. This rotation causes a further ~ 1.5 Å translation of the αM1-M2 segment towards the cytoplasmic side and a further unwinding of the αM2 cytoplasmic end between Glu144 and Ile150. Stabilizing the fully occluded state, the segment Ile150-Lys155 rewinds to form a hydrophobic cluster (24,25) that ensures tight association between αM2-A and the A-αM3 linker segment with the P domain (Fig. 7G,H).

Interestingly, similar sequential rearrangements are seen for the SERCA1a E2-BeF_x_ to E2-MgF_x_ transition when comparing thapsigargin-free and thapsigargin-bound (and proton-occluded) SERCA1a E2-BeF_x_ structures (PDB ID 3B9B and 2ZBE vs. 2ZBF (14,23)) (Figs. S4, S5). This indicates that ion transporting P2-type ATPases undergo similar conformational changes in dephosphorylation, yet differing from e.g. P1B-ATPases (26) and P4-ATPases (27,28).

#### Changes in the α-C-terminal region and β-subunit

The α-C-terminal region plays an important role in the transport cycle (6,29,30). Mutations in the region are associated with neurological disorders and the C-terminus seems integral to the function of the Na^+^ site III and protonation of K^+^-bound states (29). However, when comparing the E2-BeF_x_ and [Rb_2_]E2-MgF_x_ structures in a local αM8 helix superimposition, only subtle changes are seen. Although they may affect the position and dynamics of the conserved C-terminal tyrosine residue (Fig. S6A) and thereby solvent access and protonation of the Na^+^ site III, functionally important features are difficult to qualify from the current study. Similarly, the β-subunit TM helix (β-TM) makes a small, lateral shift towards αM7 (~2°) at Rb^+^ occlusion.

On the other hand, the flexible N-terminal tail of the β-subunit (Phe15-Ser31 modelled) undergoes a different twist in the Rb^+^ occluded state, changing its interaction with the α-subunit. In the E2-BeF_x_ state, βArg27 likely forms a salt bridge with Glu840 (αM7), which gets disrupted in the Rb^+^ occluded state, where βArg27 is instead exposed to the cytosol. The N-terminal tail of the β subunit appears more extended, as βAsn18 (the resolved β-N-terminus of [Rb_2_]E2-MgF_x_) is shifted 6.5 Å compared to E2-BeF_x_, in response to changes in position of the cytoplasmic domains of the α-subunit in the E2P-E2·P_i_ transition (Fig. S6B).

##### Binding of ions and nucleotides induce dissociation of BeF_x_ from the Na^+^,K^+^-ATPase

Monitoring the recovery of Na^+^,K^+^-ATPase activity, we investigated the effect of ions and nucleotides on the E2-BeF_x_ complex (Fig. 8). The time course of the reactivation of ATPase activity reflects the rate of BeF_x_ dissociation from the enzyme. Spontaneous dissociation in the media with 100 mM NMG^+^ (aimed to compensate for possible effects of ionic strength) is slow but increases in the presence of specific ions (Fig. 8A). K^+^ seems to be more efficient than Na^+^, as would be expected for extracellular ion binding. Note that Na^+^-concentration in the experiment is 2-fold higher than that of K^+^ while total ion concentrations are always 100 mM, adjusted with NMG^+^.

**Fig. 8.**
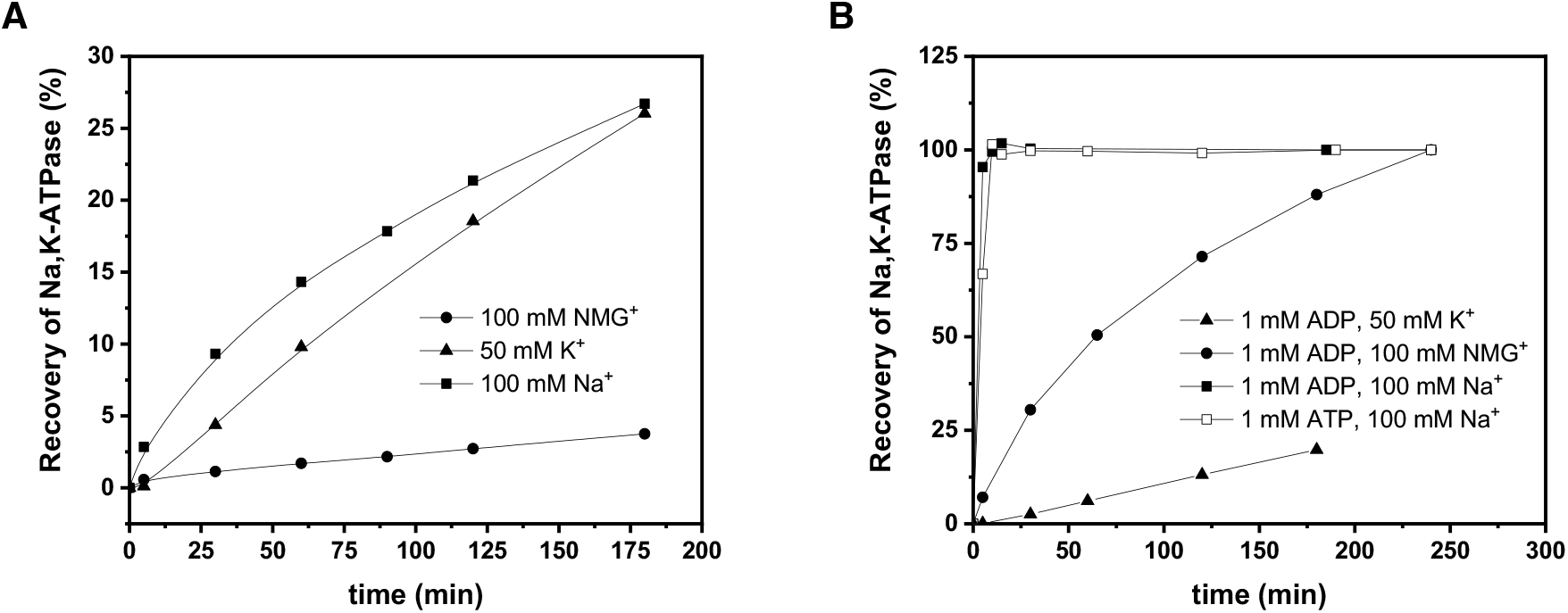
Restoration of the Na^+^,K^+^-ATPase activity due to dissociation of BeF_x_ induced by binding of cations (A) or nucleotides (B). Note that the Na^+^-concentration is two-fold higher than that of K^+^ with total ion concentration adjusted to 100 mM with NMG^+^.

Also ADP (and ATP, but ADP formation during preincubation with ATP cannot be avoided) speeded up reactivation of the ATPase activity compared to NMG^+^ alone, and this effect is further amplified by Na^+^. On the other hand, K^+^ completely cancels the ADP-effect, presumably by preventing its binding by allosterism. We attempted crystal soaking experiments with ADP and Na^+^, but saw no binding (data not shown). Presumably, ADP and Na^+^ binding would entail conformational changes that are inhibited by the crystal packing (see below).

Interestingly, the rate of dephosphorylation of the P_i_-induced E2P form does not increase upon addition of neither ADP nor K^+^ (21,31), so our data point to an apparently significant difference between the two otherwise analogous conformations (P_i_-induced E2P and E2-BeF_x_). The explanation is, however, relatively straight forward: high stability of the E2-BeF_x_ form allows both ions and nucleotides to equilibrate with the protein and to influence and stimulate BeF_x_ dissociation. The acyl-phosphate bond on the other hand is reactive, and P_i_-induced E2P dephosphorylates with the rate constant of approximately 60 min^−1^ (21), i.e. faster than equilibration with other ligands.

##### Docking of ADP into E2-BeF_x_ structure

Visualization of ADP interaction with E2-BeF_x_ was approached by docking the ADP molecule from the [Na_3_]E1-AlF_x_-ADP structure (PDB ID 3WGV(32)) into the E2-BeF_x_ state guided by a local structural alignment of the N-domains. ADP fits well into the N domain of the E2-BeF_x_ complex (with Asp443 and Glu446 within coordination distance) and requires only minor rearrangement for the Phe475-Gln482 loop (Fig. 9A). The A domain comes into close contact with Lys205 near the β-phosphate of ADP (< 1.9 Å), which presumably will cause a slight repositioning of the A domain. In the E2-MgF_x_ state, the interface between the N and A domains is only stabilized by an ionic bond between Arg544 (N domain) and Glu216 (A domain). Thus, it is likely that Arg544 either interacts with the β-phosphate of ADP upon its binding to E2-BeF_x_ (and allows further interaction with Na^+^) or forms a salt bridge with Glu216 as consequence of interactions with K^+.^. Both destabilizing effects are likely to provide greater mobility of the A domain and accelerate BeF_x_ dissociation.

**Fig. 9.**
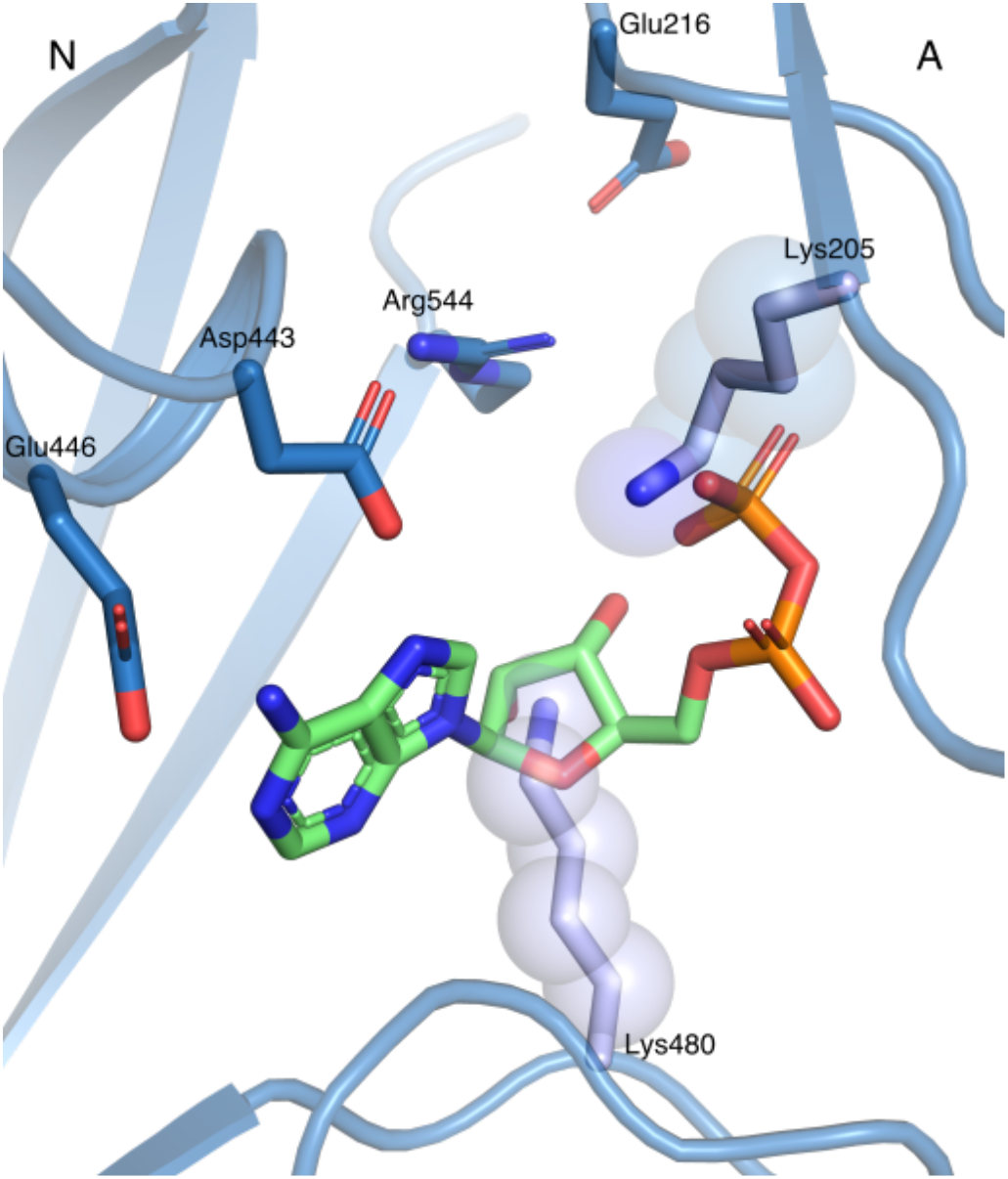
ADP from the [Na_3_]E1-AlF_4_^−^-ADP structure (PDB ID 3WGV(32)) modelled into the E2-BeF_x_ complex. N-domain alignment of the E2-BeF_x_ and the [Na_3_]E1-AlF_4_^−^-ADP structures showed a good fit for ADP at the nucleotide site with only few readjustment needed.

## Discussion

Comparison of the functional properties of the P_i_-induced E2P and E2-BeF_x_ revealed close similarities of these forms: kinetics of interactions with the cardiotonic steroids and K^+^ are virtually the same, and the structures of E2-BeF_x_ and E2P stabilized by cardiotonic steroids show clear structural resemblance.

The cation binding sites are open to the extracellular side and accept Na^+^, K^+^ as well as Mg^2+^ at low affinity. The ability of E2-BeF_x_ to occlude Na^+^ is lost, even in the presence of oligomycin, and Na^+^ affinity is therefore low. Na^+^ binding, however, destabilizes E2-BeF_x_, in a way analogous to the dephosphorylation under the Na^+^/Na^+^ exchange reaction associated with ATP hydrolysis (33).

Binding of K^+^ (or the congener Rb^+^) triggers the conformational re-arrangements necessary for ion occlusion and destabilization of the Asp369-BeF_x_ bond. Rb^+^ soaking of the E2-BeF_x_ crystals revealed a sequence of Rb^+^ pre-occlusion steps. Although only qualitative, the occupancy analysis revealed that the initial (Rb)E2-BeF_x_ binding form has only partial occupation at site I. Further, the anomalous difference suggests an overlap of the initial site with the Mg^2+^ site (Fig. 6B). This indicates, that going from the E2-BeF_x_ open state, K^+^ will bind in a sequential manner by first binding to an initial site that likely overlaps with the low-affinty Mg^2+^ site coordinated by Asn776, Glu779 (αM5) and Asp804 (αM6) (10). Subsequenty, the initial site changes configuration to the K^+^ site I, at the same time making space for ion binding at site II. That involves mainchain carbonyls of αM4 and ultimately leads to closure of the extracellular gate. This mechanism, described from the outward-open E2-BeF_x_ form and onwards, is consistent with exchange kinetics followed in the opposite direction. Thus, in the occluded [K2]E2-MgF_x_ form, the exchange with extracellular ions occurs first at site II (34). The oscillations of αM3-M4 helices allowing only one ion to bind at the time stand for the “flickering-gate”, first proposed by Forbush (35) on the basis of deocclusion experiments and later supported by high-speed voltage clamp measurements (15).

In summary, the conformational transitions observed in soaking experiments are the prelude to the dephosphorylation process in the Na,K-ATPase reaction. The sequential binding of two K^+^ through the αM1-M2-triggered rotation of the A domain re-arranges the TGES motif, allowing water access and desphosphorylation (Fig.S7).

The above interactions with K^+^ suggest that the E2-BeF_x_ complex is a mimic of the ground state E2P. However, E2-BeF_x_ is also destabilized by ADP, i.e. it has a property of an ADP-sensitive state (Fig.8B). The ability of ADP to induce a direct hydrolysis of the acyl-phosphate bond was described earlier by Hobbs et al. (4) for the phosphoenzyme E*P. ADP-induced dephosphorylation of E*P occurred directly, without transition to E1P. In the case of E2-BeF_x_, however, a transition towards E1-BeF_x_ might be induced by the simultaneous presence of ADP and Na^+^, since the synergy between these two ligands is very clear (Fig. 8B). K+ on the other hand, cancels the ADP-effect (Fig. 8B). Due to occlusion, it prohibits ADP binding and leads to slow BeFx dissociation from the [K_2_]E2-BeF_x_ complex.

In conclusion, the E2-BeF_x_ complex of Na^+^,K^+^-ATPase represents a phosphorylated intermediate that can reach both E2P and E*P. The structural similarity to the backdoor phosphorylated (P_i_-induced) E2P form is obvious from its comparison with the CTS-stabilized complexes of the Na,K-ATPase. Functionally, P_i_-induced E2P and E2-BeF_x_ react nearly identically with CTS (here demonstrated by anthroylouabain interactions) and with K^+^(probed by RH421 fluorescence). Both forms are capable of Rb^+^ or K^+^ occlusion. Soaking of the E2-BeF_x_ crystals with Rb^+^ allowed visualization of different steps in the occlusion process. At the same time, firm coordination of BeF_x_ within the phosphorylation site granted time for nucleotide binding and exploration of a dynamic E*P intermediate. These studies should be very informative for future investigations of P-type ATPase dynamics and studies of e.g. mutant forms.

## Experimental procedures

### Protein preparation

Pig kidney Na^+^,K^+^-ATPase was purified as previously described (36). The specific ATPase activity of the Na^+^,K^+^-ATPase purified membrane preparations was about 1,800 μmol P_i_ per hour per mg of membrane protein at 37 °C.

### Crystallization and data collection

The BeF_x_ inhibited complex of Na^+^,K^+^-ATPase was formed by pre-incubation of the membraneous enzyme in 20 mM histidine pH 7.0, 10 mM NaF, 0.5 mM MgCl_2_, 20 μM BeSO_4_. The stabilized E2-BeF_x_ complex was subsequently solubilized in the same buffer with the non-ionic detergent octaethyleneglycol mono-n-dodecylether [C_12_E_8_] at a ratio of 0.9 mg C_12_E_8_ per mg protein, and insoluble material was removed by ultracentrifugation. The final concentration of solubilized protein was 9-10 mg/mL.

Crystals were grown by vapour diffusion at 15 °C in 2 μl hanging drops for 2-3 weeks. The protein sample was mixed in a 1:1 ratio with reservoir solution containing 16.5 % (wt/vol) polyethylene glycol 2000 monomethyl ether (PEG 2000 MME), 10 % (vol/vol) glycerol, 175 mM MgCl_2_, 150 mM NaCl, 20 mM HEPES/MES pH 6.2 and 2 mM DTT. The crystals were dehydrated overnight at 4 °C against a 30 % PEG 2000 MME reservoir solution before flash-cooling. The final datasets were collected at 100 K on the DESY-EMBL beamline P14 and the SLS-X06DA (PXIII) beamline. For Rb^+^ soaks, 1.1 mM sucrose monodecanoate was added to the solubilized protein before mixing with reservoir solution. 10 mM and 50 mM RbCl, respectively, was added to the crystallization drop and equilibrated for 20 seconds and 3 hours, respectively. The final datasets of Rb^+^ soaked crystals were collected on the DESY-EMBL beamline P13 and the DLS-I24 beamline using a wavelength of 0.814 Å.

### Data processing

Datasets were processed and scaled using XDS software (37). The crystals showed P2_1_2_1_2_1_ space group symmetry with unit cell dimensions a = 116.3 Å, b = 117.8 Å and c = 490.9 Å and two αβγ-heterotrimers per asymmetric unit. For the native E2-BeF_x_ structure, datasets derived from two crystals were merged to yield the final dataset. The data were anisotropcially scaled using the Diffraction Anisotropy Server (http://services.mbi.ucla.edu/anisoscale) (38), setting the resolution limits to 5.4 Å, 4.4 Å and 4.0 Å along a*, b* and c*, respectively. Molecular replacement was performed using PHASER (39). As serach models, we used the crystal structure of Na^+^,K^+^-ATPase E2P ouabain complex (PDB ID 4HYT (10) and the structure of a different crystal form of the E2-BeF_x_ complex (PDB ID 7D91) (9). Rigid body refinement followed by simulated annealing refinement protocol was performed in PHENIX (40). Manual refinement was carried out in Coot (41), and the further model refinement was continued in PHENIX using non-crystallographic symmetry (NCS), translation-libration-screw (TLS) parameterization, and grouped atomic displacement parameter (ADP) refinement. Because of low resolution of the data, tight geometry restraints were imposed on the model to stabilize the refinement. Rigid body groups were defined by the A, N, and P domains along with the αM1–2, αM3–4, αM5–10/ βM/γM, and β-ectodomain. NCS and TLS groups were defined by the A, N, and P domains, the transmembrane domain αM1–10/βM/γM, and the β-ectodomain. The quality of the final model was assessed using the MOLPROBITY server (42). For Rb^+^-soaked data sets, the diffraction data were first scaled using Scaleit (43) to the Na^+^,K^+^-ATPase E2P ouabain complex (PDB ID 4HYT)(10), and the initial phases were obtained by molecular replacement using PHASER and the E2-BeF_x_ structure. Rigid body refinement and NCS refinement (same groups as for the E2-BeF_x_ structure) were performed for each data set. Occupancy calculations were performed using molecular replacement with single-wavelength anomalous diffraction (MR-SAD (44)). All protein structure figures were prepared using PyMOL (The PyMOL Molecular Graphics System, Version 2.3.0 (Schrödinger LLC, 2012)).

### Na^+^ -occlusion

The membrane-bound enzyme (0.5 mg/ml) was incubated with 1 mM ^22^NaF in the presence of 10 mM Tris-HCl pH 7.0, 0.5 mM MgCl_2_, 1 mM ^3^H-glucose in the presence or absence of 0.02 mM BeSO_4_, with or without 0.02 mg/ml oligomycin for 5 hours at 0°C. Then, 0.4 ml of the mixture was loaded on a Millipore HAWP 0.45 μm filter and immediately washed with 1 ml of Tris-HCl buffer containing 20 mM NaCl to decrease background. The filters were counted in 4 ml Packard Filtercount scintillation fluid. ^3^H-counts gave an estimate for the non-specific absorption of the isotopes of the filter and for the amount trapped in the wetting volume.

### Dissociation of BeF_x_ from the Na^+^,K^+^-ATPase induced by different ligands

BeF_x_-complex of Na^+^,K^+^-ATPase was formed by pre-incubation of the enzyme in 20 mM histidine pH 7.0, 10 mM NaF, 0.5 mM MgCl_2_, 20 μM BeSO_4_ on ice overnight. The suspension was subjected to centrifugation 30 min × 50.000g at 4 °C, and the pellet was resuspended in the media containing 20 mM histidine pH 7.0 and varying ligands in concentrations as noted in Fig.8. After different periods of incubation at 37°C, the aliquots from each sample were used for measurement of the Na^+^,K^+^-ATPase activity under optimal conditions (45).

#### Fluorescence spectroscopy

##### RH421 fluorescence experiments

Equilibrium and transient RH421 fluorescence in response to varying concentrations of KCl was measured at room temperature in 20 mM histidine pH 7.0, 10 mM NaF, 0.5 mM MgCl_2_, 0.05 mg/ml enzyme, 0.2 μg/ml RH421. Formation of BeF_x_ complex was ensured by addition of 20 μM BeSO_4_. Measurements under equilibrium conditions were performed on a SPEX Fluorolog fluorometer in a cuvette with 1 cm lightpath with continuous stirring. Excitation wavelength was 580 nm (slit 2 nm), emission 630 nm (slit 14 nm). Observed rate constants were obtained in the experiments using rapid-mixing stopped-flow spectrofluorometer (Applied Photophysics) at excitation wavelength 580 nm (slit 1 nm) with 630 nm cutoff filter on the emission side.

##### Anthroylouabain fluorescence experiments

Equilibrium and transient experiments with anthroylouabain were performed at room temperature in either 20 mM Histidin pH 7.0, 4 mM H_3_PO_4_ (adjusted with N-methyl-D-glutamine), 4 mM MgCl_2_, 0.05 mg/ml enzyme (P_i_-induced E2P complex); or 20 mM Histidin pH 7.0, 10 mM NaF, 0.5 mM MgCl_2_, 20 μM BeSO_4_, 0.05 mg/ml enzyme (BeF_x_-complex). When necessary, K^+^ was added as KCl in the experiments with P_i_-induced E2P complex form, while in case of E2-BeF_x_ complex, 10 mM NaF was replaced by equimolar KF. A SPEX Fluorolog fluorometer was used to monitor slow reactions, excitation at 370 nm (slit 5 nm), emission 485 nm (slit 5 nm). Association rate constants were obtained in the experiments using rapid-mixing stopped-flow spectrofluorometer at excitation wavelength 370 nm (slit 1 nm) with 485 nm cutoff filter on the emission side.

Dissociation of anthroylouabain complexes was performed in a chase experiment as follows: the enzyme complexes were pre-incubated with 0.2 μM anthroylouabain overnight. Then 3.3 mM ouabain was added to the sample (anthroyloaubain was diluted to 0.13 μM simulatneously) and decrease in fluorescence was monitored in time.

##### Two-electrode voltage-clamping

The experiments were performed as described in (46) In brief, mRNA was made with the mMESSAGE mMACHINE T7 Ultra Kit (Life Technologies) from linearized DNA encoding the human α2 with mutations to decrease ouabain sensitivity (Q116R and N127D) and β1. *Xenopus laevis* oocytes (EcoCyte Bioscience) were injected with 9.5/2.5 ng α2/β1 subunit mRNA in 50.6 nL. Oocytes were incubated 2-8 days at 12-18 degrees Celcius in ND96 buffer supplemented with 25 μg/ml gentamicin and 2.5 mM sodiumm pyruvate. Oocytes were clamped with an OC-725C amplifier (Warner Instruments), and the signal was digitized by a 1440A digitizer (Molecular Devices). Data was recorded with PClamp 10.4 (Molecular Devices) and analysed with ClampFit (Molecular Devices) and GraphPad Prism (GraphPad Software). For sodium/sodium exchange recordings, the extracellular buffer contained 115 mM NaOH, 110 mM sulfamic acid, 10 mM Hepes, 5 mM BaCl_2_, 1 mM MgCl_2_, 0.5 mM CaCl_2_, 10 μM ouabain, pH 7.4 (adjusted with sulfamic acid). For other recordings, NaOH was replaced by NMDG or KOH as indicated. Voltage jumps were performed from a holding potential of −50 mV with 200 ms jumps to potentials between 60 mV and −160 mV in 20 mV steps. Charge (Q) and rate currents were determined by subtracting a recording with 10 mM ouabain added from the immediately preceding recording. On and off-currents were fitted with single exponentials to determine rate currents and charge, respectively. Charges were fitted to a Boltzmann distribution. For steady state currents, the voltage protocol was performed in buffer, buffer with 10 mM ouabain, and buffer. The last recording ensures stability throughout the recordings.

## Data availability

Coordinates and diffraction data for the E2-BeF_x_ structure and for the low resolution models of the initial, early and late stage of Rb+ binding have been deposited in the Protein Data Bank under accession code 7QTV for the refined E2-BeFx structure, and XXXX, and YYYY for the low resolution models of the Rb+ binding forms of E2-BeFx.

## Supporting information

This article contains supporting information.

## Author contributions

**Marlene U. Fruergaard**: Formal analysis, Investigation, Writing – Original Draft, Writing – Review and Editing, Vizualization; **Ingrid Dach**: Formal analysis, Investigation, **Jacob L. Andersen**: Verification, Data curation; **Mette Ozol**: Formal analysis, Investigation, Vizualization; **Azadeh Shasavar:** Formal analysis, Verification, **Esben M. Quistgaard**: Formal analysis, Verification; **Hanne Poulsen**: Verification, Funding acquisition; **Natalya U. Fedosova:** Conceptualization, Methodology, Supervision, Investigation, Formal analysis, Writing – Original Draft, Writing – Review and Editing, Funding acquisition; **Poul Nissen**: Conceptualization, Methodology, Supervision, Verification, Writing – Original Draft, Writing – Review and Editing, Project administration, Funding acquisition

## Acknowledgments

This work was supported by grants from the Danish National Reserarch Foundation through the Pumpkin center of excellence (to P.N.), the Danish Council for Independent Research (DFF-7016-00125, to N.U.F.), the A.P. Møller Foundation for the Advancement of Medical Science (to N.U.F.), the Lundbeck Foundation (R155-2015-2666 and R310-2018-3713 to P.N., Rxxx-201X-xxxx to H.P.). We are grateful to B. Bjerring Jensen, Tetyana Klymchuk and Anna Marie Nielsen for excellent technical assistance, and to Jesper Karlsen for support with scientific computing. We thank Tomas Heger and Samuel Hjorth-Jensen for screening Na^+^ and ADP soaked crystals..

## Supporting Information

**Fig. S1.**
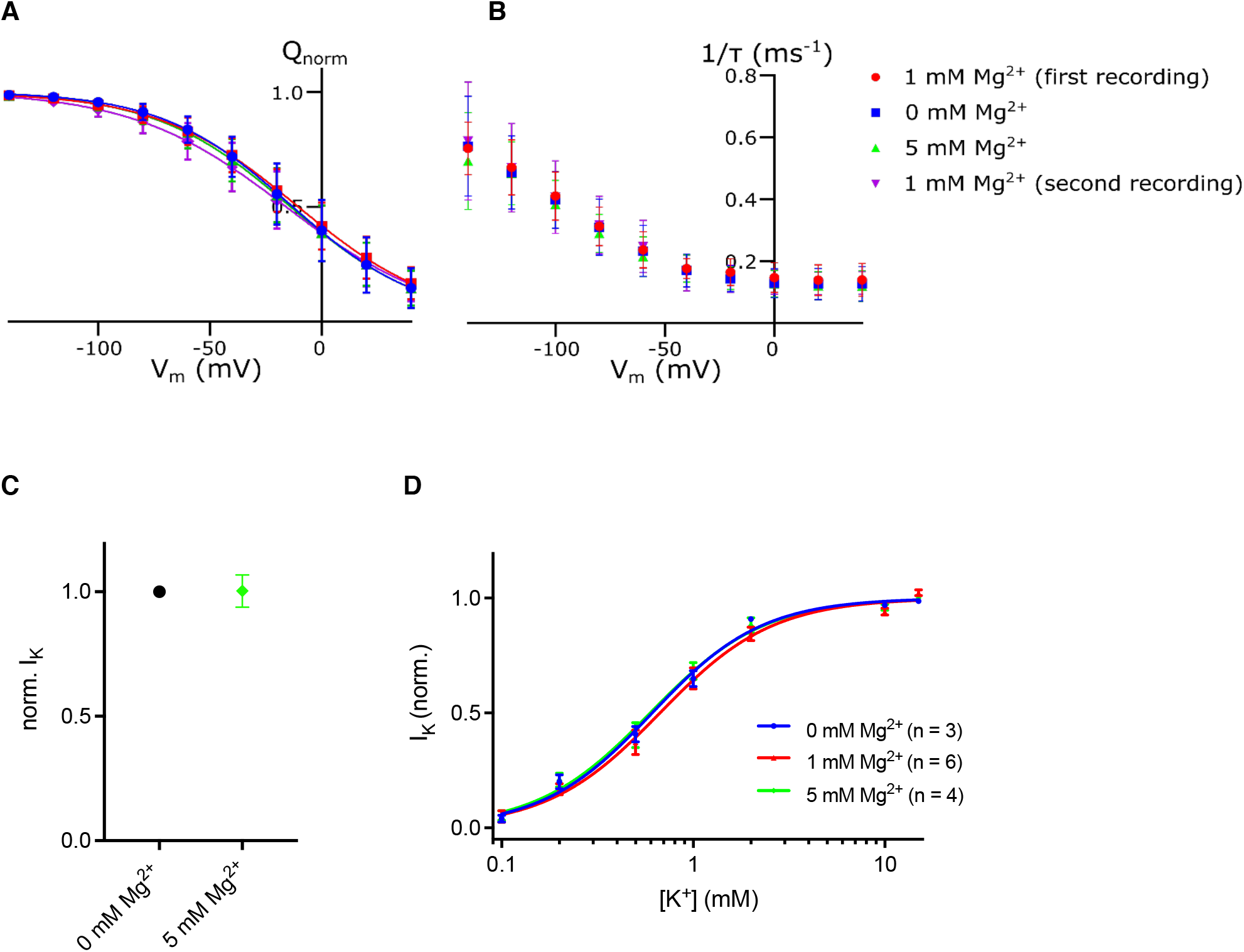
Magnesium effect on sodium and potassium binding. A) Voltage jumps under sodium/sodium exchange conditions in oocytes expressing human α2β1 (n=7). The ouabain-sensitive currents are determined at three magnesium ion concentrations for each oocyte in the order 1 mM, 0 mM, 5 mM, 1 mM to ensure that the measurements were stable. Q/Vm curves and B) rate constants were determined for each concentration of magnesium. C) The efficacy of potassium was tested at −50 mM and 100 mM potassium as the pumping current (I) in the presence and absence of 5 mM magnesium (n=4). D) Potassium affinity was determined with 0 mM, 1 mM and 5 mM magnesium.

**Fig. S2.**
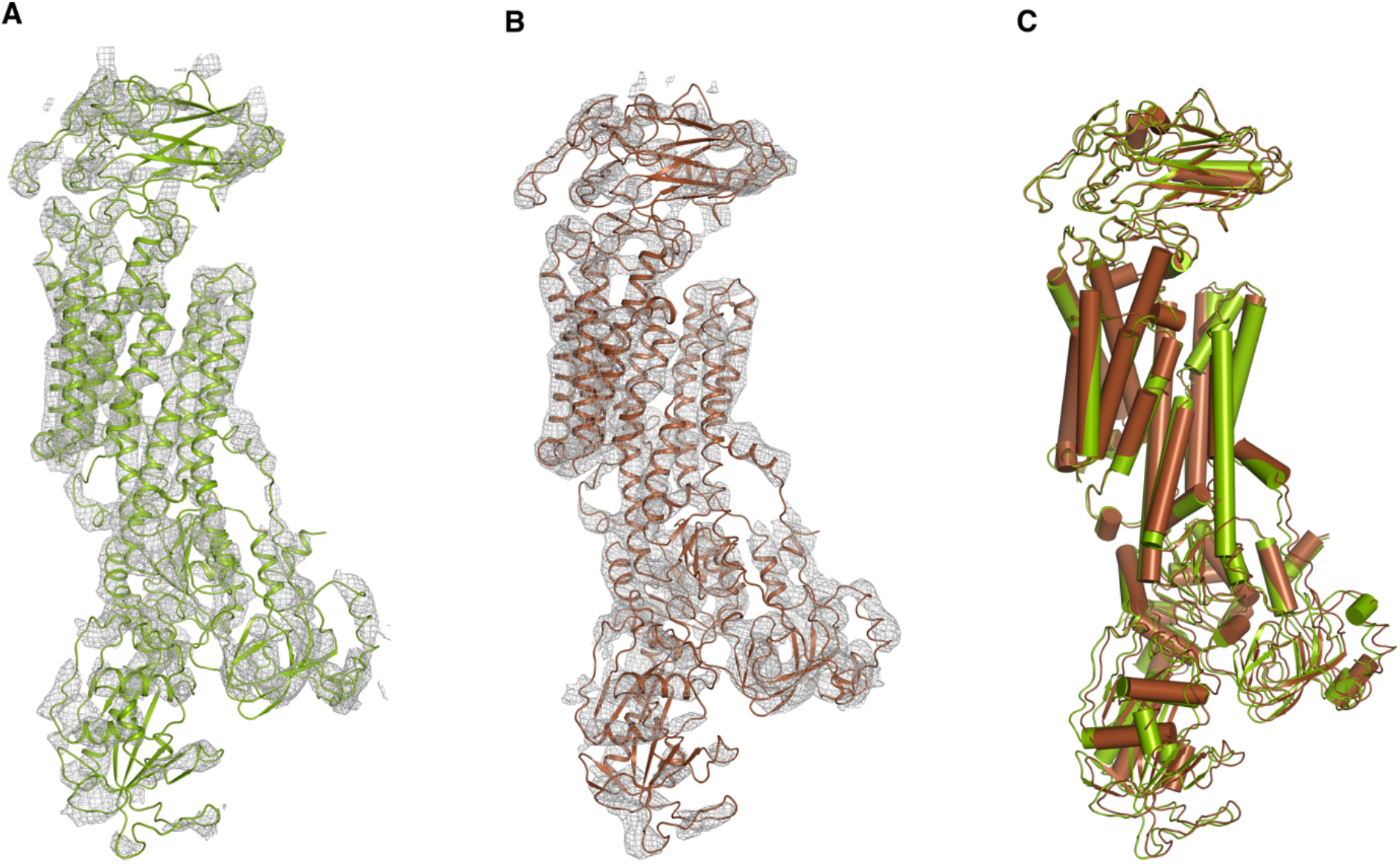
(A,B) Electron density maps for two protomers of the (Rb)E2-BeF_x_ initial site form (10 mM Rb^+^, 20 sec). 2Fo-Fc is depicted in gray mesh (contoured at 1.3 σ). (C) αM7-M10 transmembrane alignment of the two protomers showing high resemblance (root mean square deviations (rmsd) = 1.16 Å, all C_α_).

**Fig. S3.**
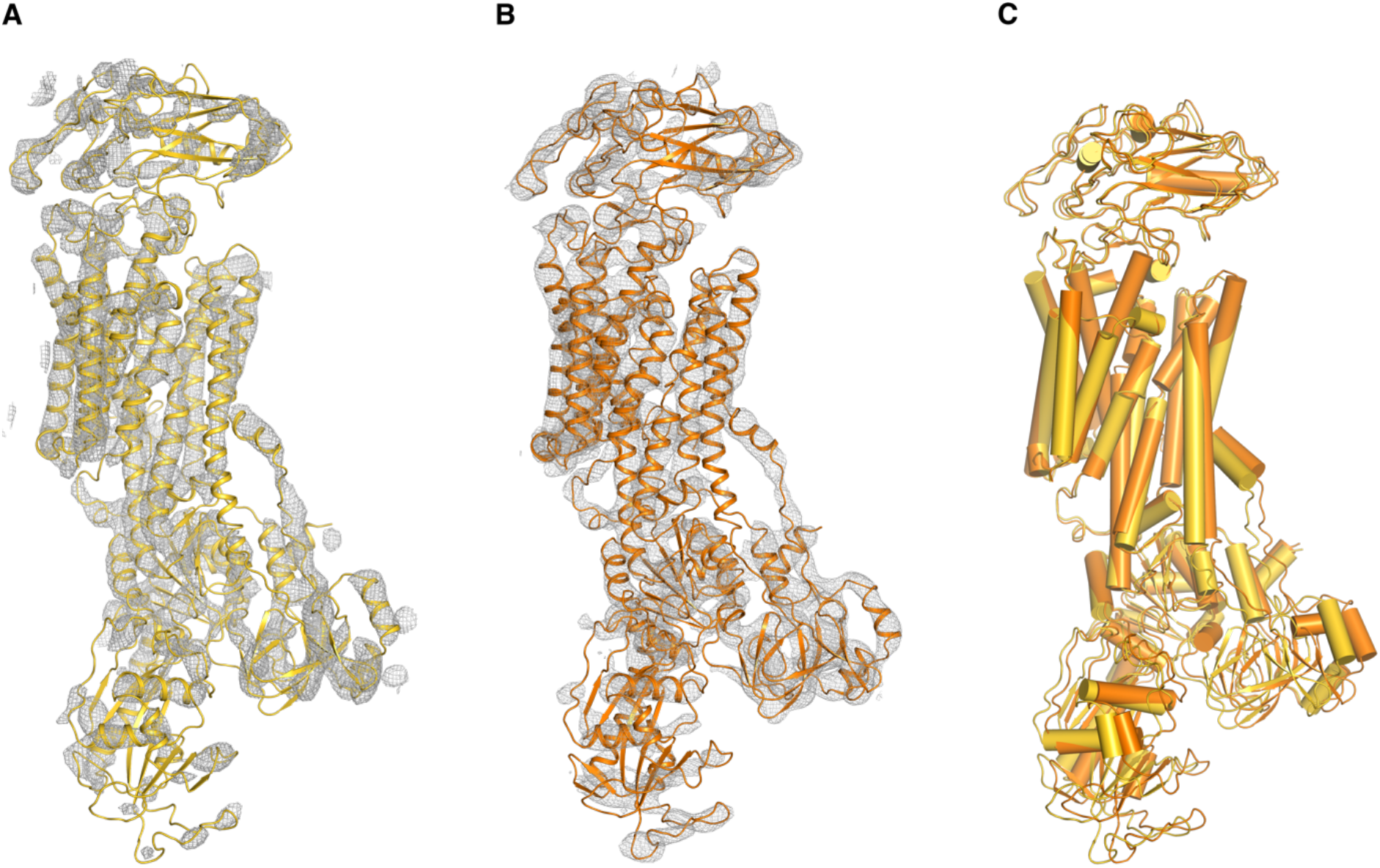
Electron density maps for (A) the early (Rb)E2-BeF_x_ form (protomer 1) and (B) the late (Rb)E2-BeF_x_ form (protomer 2) from the 50 mM Rb^+^-soaked crystal (50 mM Rb^+^, 3 hour). 2Fo-Fc is depicted in gray mesh (contoured at 1.3 σ). (C) αM7-M10 transmembrane alignment of the two protomers (rmsd = 1.96 Å, C_α_).

**Fig. S4.**
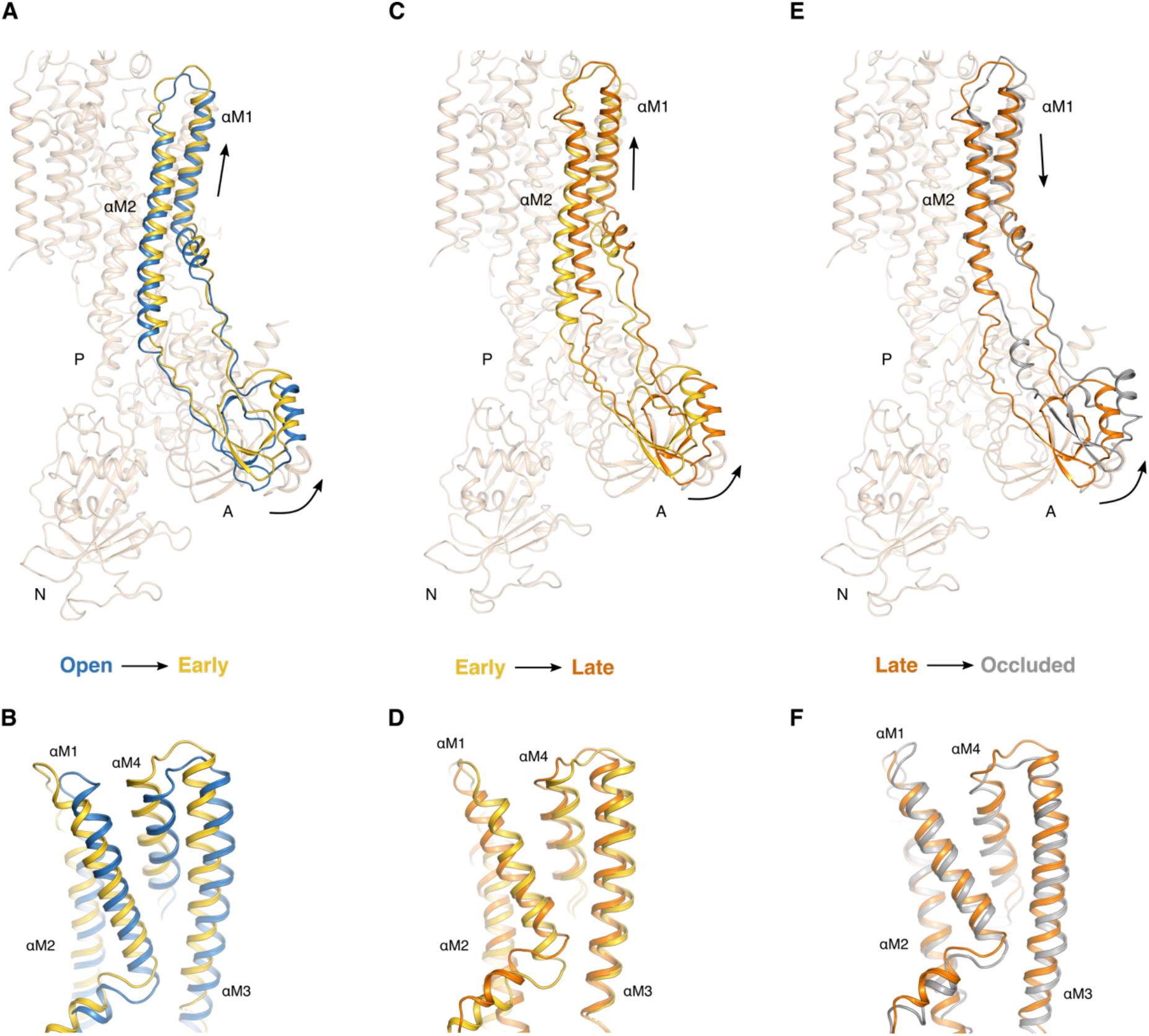
Sequential structural changes from (A,B) the E2-BeF_x_ open state (blue) to (Rb2)E2-BeF_x_ form early (gold), (C,D) early form to (Rb2)E2-BeF_x_ late form (orange), and (E,F) late form to fully occluded [Rb_2_]E2-MgFx state (gray) (PDB code 3KDP(1)). For clarity, domains P, N, αM3-M10, β- and γ-subunits for all structures (except E2-BeF_x_ open state) have been removed in the upper panel (A,C,E). Only the regions showing major conformational re-arrangements between the states have been highlighted.

**Fig. S5.**
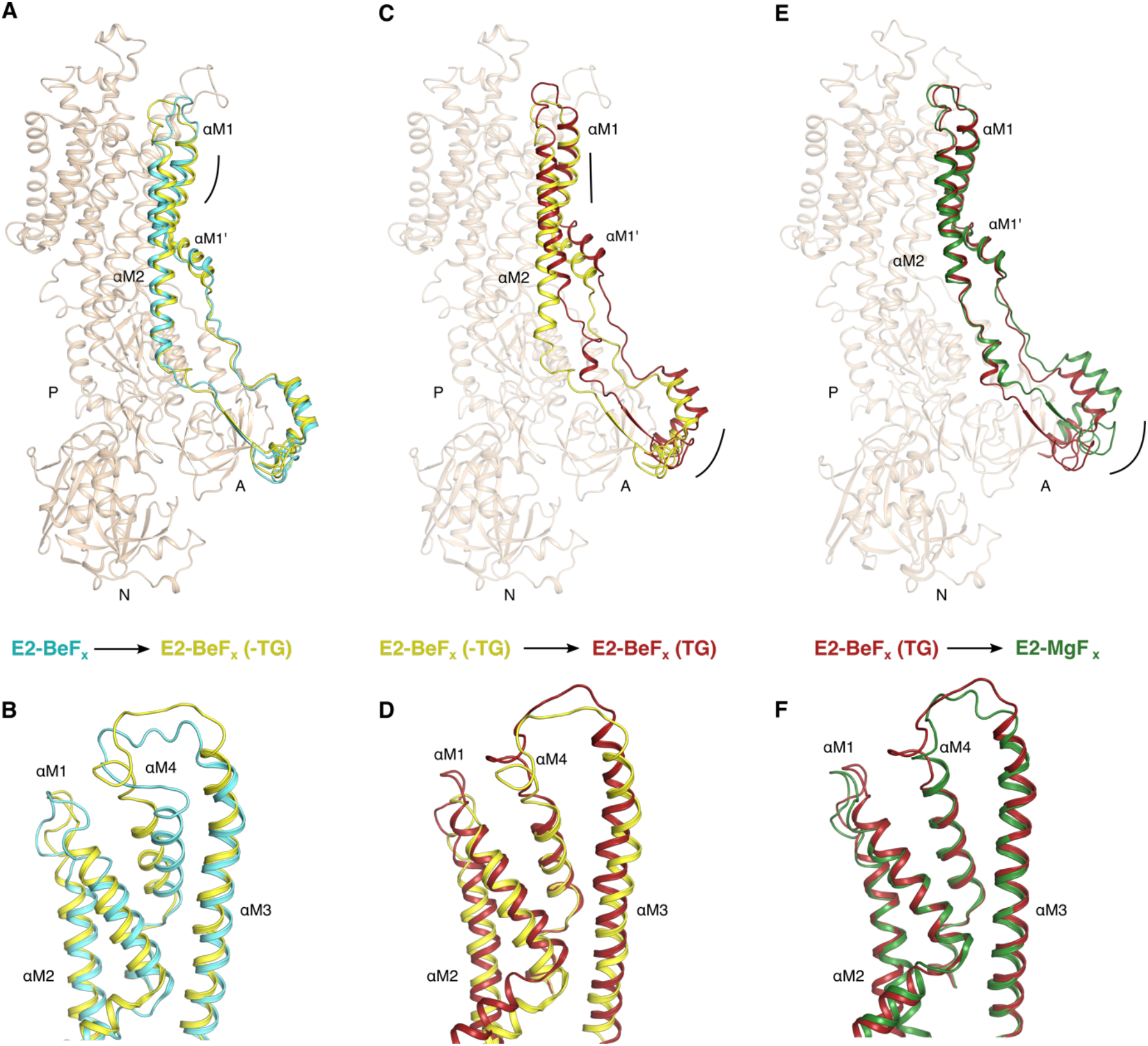
Sequential structural changes from (A,B) the SERCA E2-BeF_x_ open state (PDB ID 3B9B (2); cyan) to E2-BeF_x_ thapsigargin free-form (PDB ID 2ZBE (3); yellow), (C,D) from E2-BeF_x_ thapsigargin free-form to E2-BeF_x_ thapsigargin free-bound (PDB ID 2ZBF (3); red), and (E,F) from thapsigargin free-bound to E2-MgFx (PDB ID 3FGO (4); green). For clarity, domains P, N and αM3-M10 for all structures (except E2-BeF_x_ open state) have been removed. Only the regions showing major conformational re-arrangements between the states have been highlighted.

**Fig. S6.**
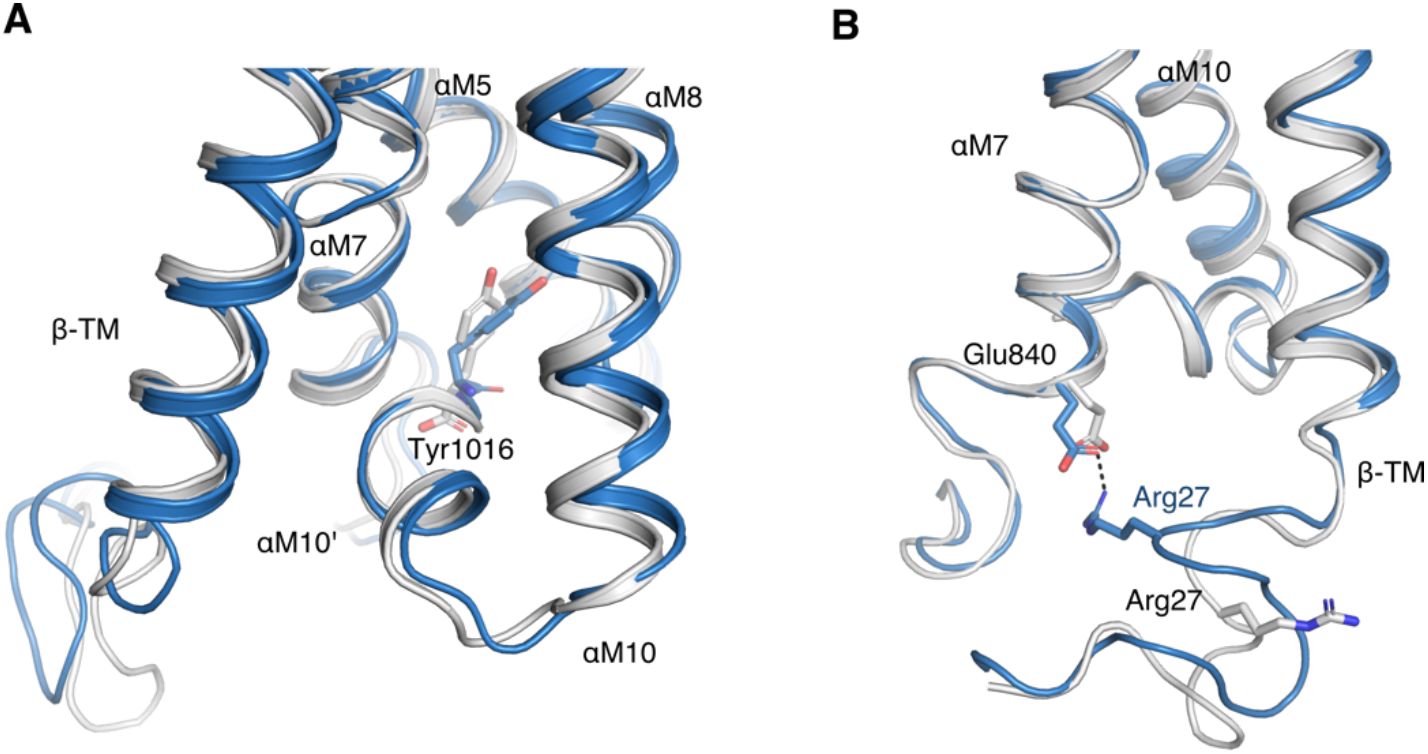
(A) Flexibility at the C-terminal Tyr1016 associated with H^+^ leak currents. Structures are here aligned based on a local αM8 superimposition. (B) Movement of the β-TM helix and its cytosolic N-terminal segment. The ionic interaction of Arg27 (β) and Glu840 (αM7) is disrupted upon Rb^+^ occlusion as the β N-terminal is changing its position due to movements of the cytoplasmic domains of the α-subunit in formation of the dephosphorylation complex. The structures were aligned as in A.

**Fig. S7.**
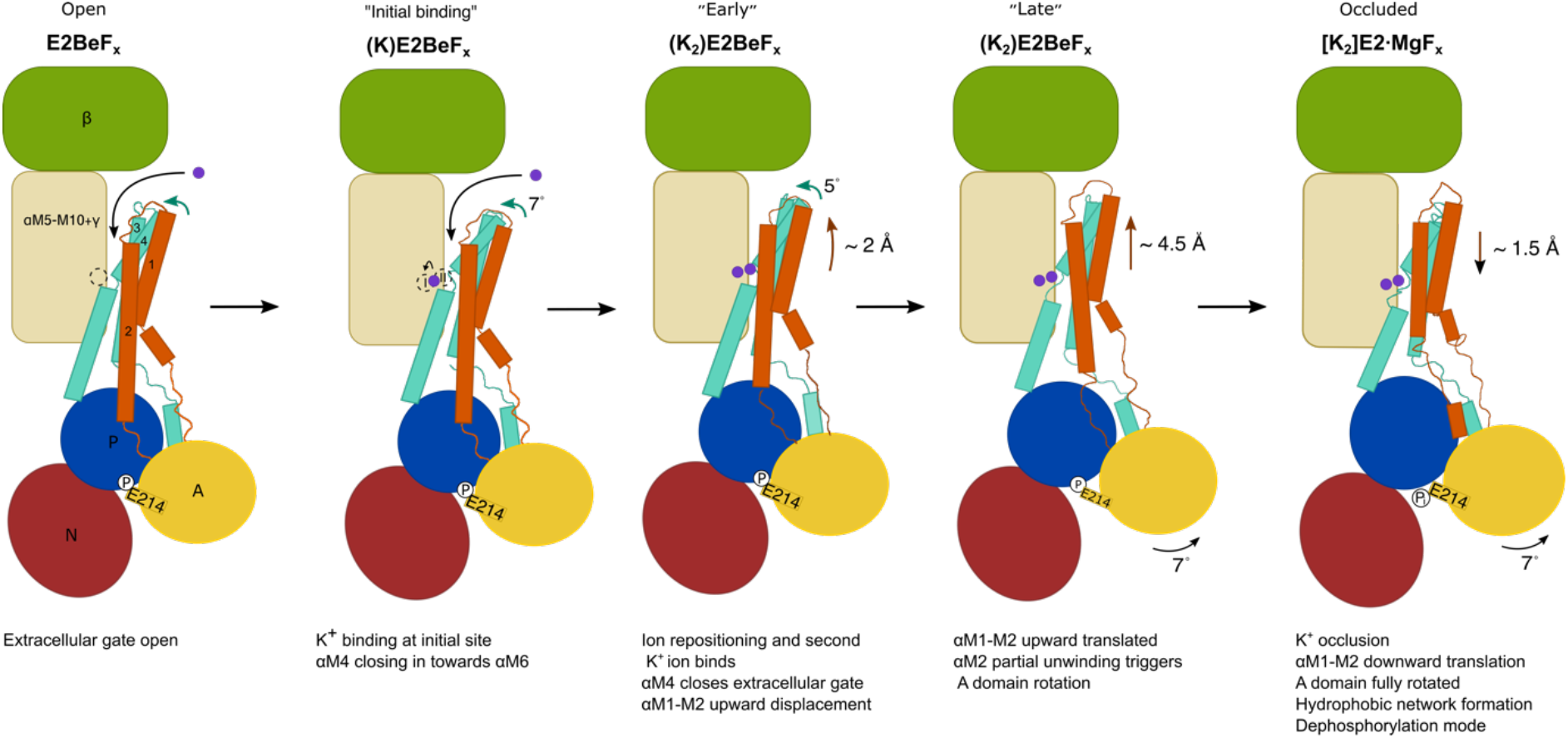
Consequences of K^+^ binding within the extracellular sites of the Na^+^,K^+^-ATPase (cartoon). The sequence of events starts with binding of one ion at the initial site (overlapping with the Mg^2+^ site), dragging αM4 towards M6. As the second ion binds, the closure of the gate, pushes the M1-M2 segment ~2 Å upwards towards the extracellular side through van der Waals contacts, which causes the A domain to rotate ~7º around the phosphorylation site (towards the membrane away from the N domain). Following a further small rotation of the A domain and αM1-M2 relocation, the dephosphorylation reaction becomes catalyzed by the TGES motif, which leads to the full occlusion of K^+^ ions ([K_2_]E2P_i_ state(1)).

**Table S1.**
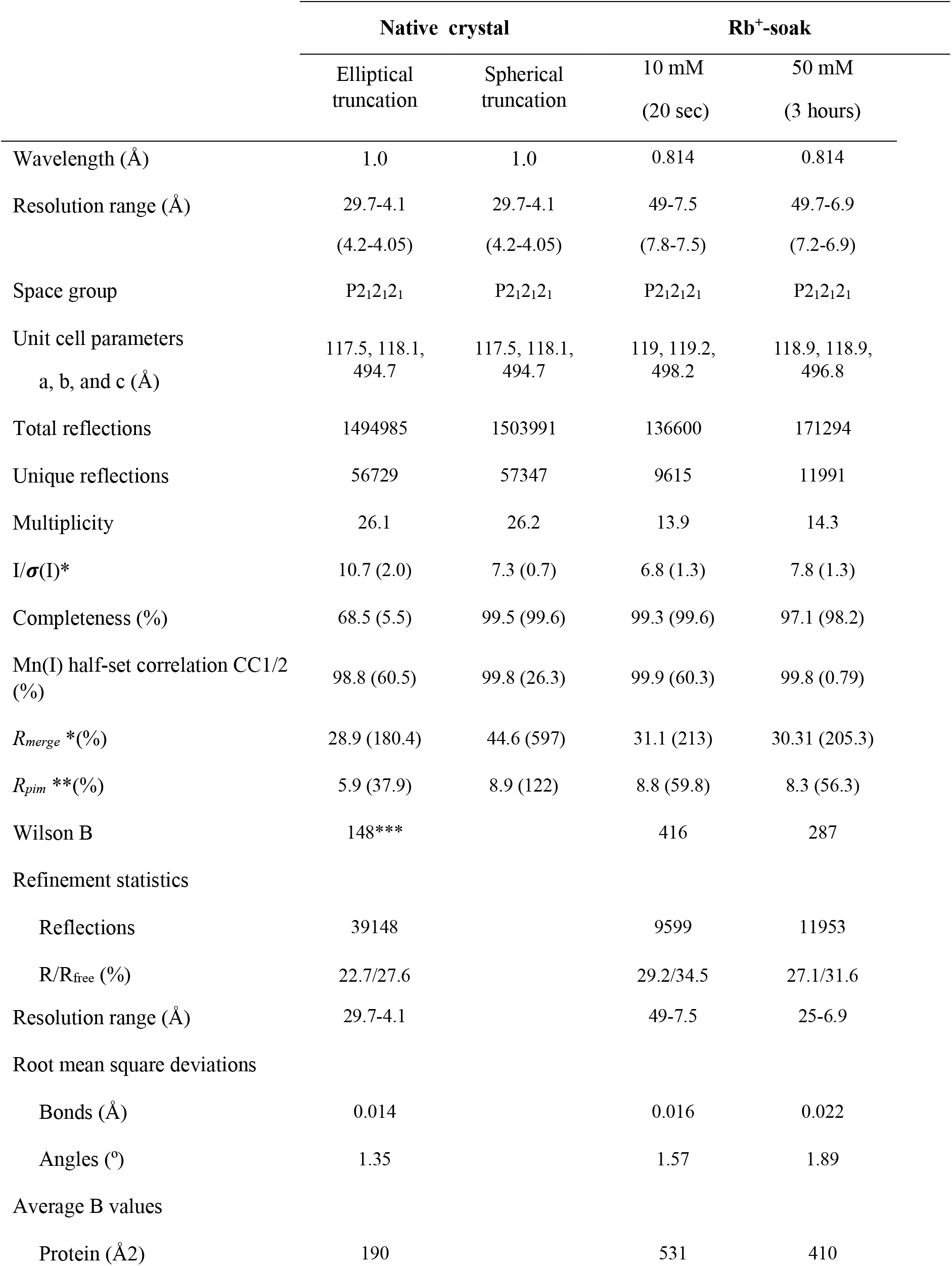
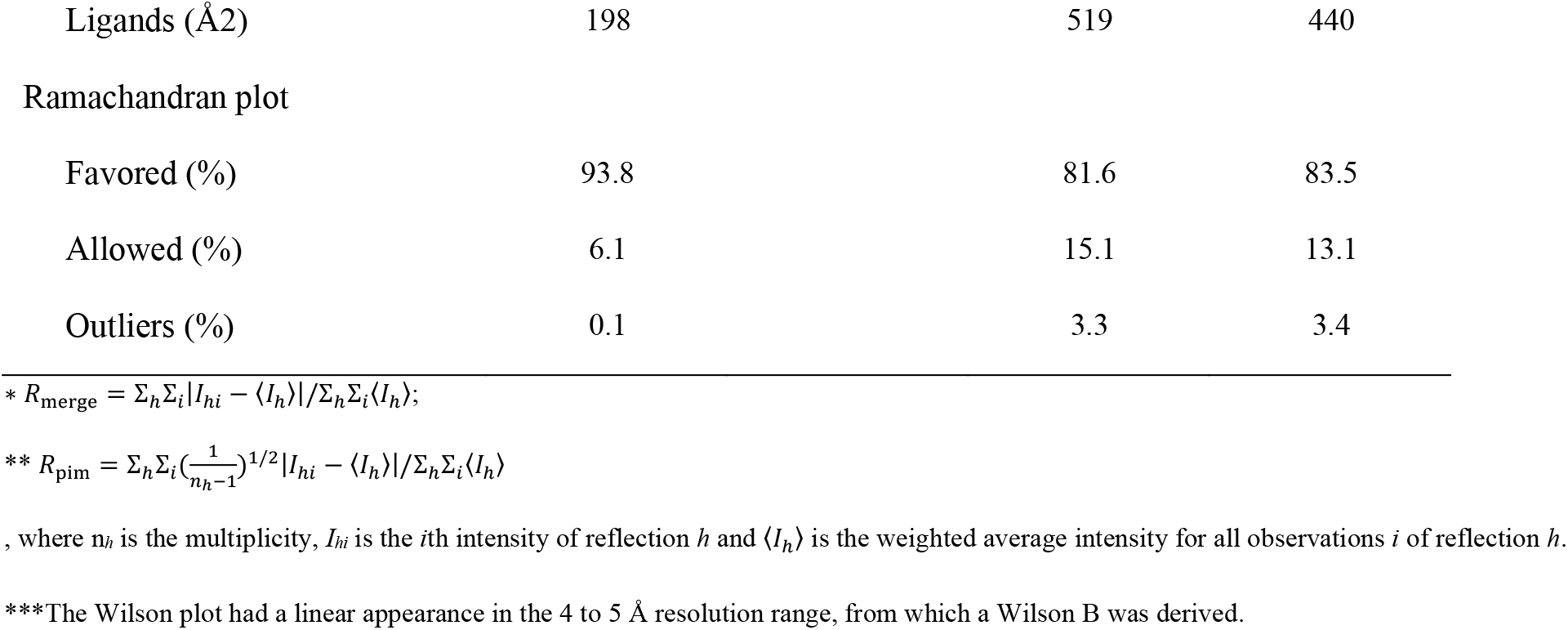
Data collection and model refinement statistics. Values in brackets are the highest resolution shell.

